# Exploring the activity and the essentiality of the putative Δ6-desaturase in the procyclic and bloodstream forms of *Trypanosoma brucei*

**DOI:** 10.1101/2023.11.23.568515

**Authors:** Michela Cerone, Terry K Smith

## Abstract

Trypanosomatids have been shown to possess an exclusive and finely regulated biosynthetic pathway for *de novo* synthesis of fatty acids (FAs) and particularly of polyunsaturated fatty acids (PUFAs). The key enzymes for the process of unsaturation are known as desaturases. In this work, we explored the biocatalytic activity of the putative Δ6-desaturase (Tb11.v5.0580) in the native organism *T. brucei*. Utilising fatty acid analysis *via* GC-MS, we were able to elucidate *via* genetic manipulation of the level of expression of Δ6-desaturases in both procyclic (PCF) and bloodstream (BSF) forms of *T. brucei* and *via* supplementation of the media with various levels of FA sources, that docosahexaenoic acid (22:6) and/or docosapentaenoic acid (22:5), and arachidonic acid (20:4) and/or docosatetraenoic acid (22:4) are the products and the substrates respectively of this Δ6-desaturases. Interestingly, we were able to observe, *via* lipidomic analysis with ESI-MS/MS, an increase in inositol-phosphoryl ceramide (IPC) in response to the overexpression of Δ6-desaturases in low-fat media, both in PCF and rather surprisingly in BSF. The formation of IPC is normally only observed in the stumpy and procyclic forms of *T. brucei*. Therefore, the expression levels of Δ6-desaturases, which varies between BSF and PCF, might be involved in the cascade(s) of metabolic events that cause these remodelling of the lipid pools and ultimately morphological changes, which are key to the transition between these life-cycle stages.

**Author summary:** *Trypanosoma brucei* is a unicellular parasite that causes human and animal African trypanosomiasis. These parasites have the special ability to make their own pool of fat molecules by assembling and modifying the fatty acid building blocks that they take up from the human and animal hosts and from the insect vector. In this study, we investigated the unknown activity of a desaturase enzyme. By modulating its activity, we showed that it can make different levels of high-value long chain polyunsaturated fatty acids (LC-PUFAs) often known as omega-6 and omega-3. If we increase or reduce the fat sources available from the outer environment, the cells respond by making more or less LC-PUFAs and by forming different type of lipids and sphingolipids for their cellular membranes. We highlighted that by tuning the level of activity of the desaturase and varying the type and amounts of fat sources available to the cells, *T. brucei* can alter their morphology. This is key for the parasites to adapt to the various environments and the nutrients therein that are often constantly changing within the host, allowing the shift between different life-stages during the complex life cycle from the insect vector to the host and back.

## Introduction

*Trypanosoma brucei* is an extracellular kinetoplastid parasite and the causative agent of Human and Animal African Trypanosomiasis (HAT, AAT).(**1–3**) *T. brucei* is transmitted by the Tsetse fly to humans and livestock animals when it takes a blood meal. In the insect vector, the parasites proliferate as procyclic form (PCF) and differentiate into epimastigotes, and finally into the infective metacyclic trypomastigotes within the salivary glands of the Tsetse fly. (**1,2**) Once in the bloodstream, the parasites transform in slender bloodstream form (BSF) and circulate in the bloodstream, but also invade the extracellular space of many organs including the skin, adipose tissue, lymph and eventually brain. (**4,5**) Some of the BSF *T. brucei* responding to a quorum sensing mechanism routinely differentiate into the pre-adaptive and non-replicative stumpy form, which is pre-adapted to survive when taken up by the fly with another blood meal. (**1,6–8**)

Throughout this complex life cycle, *T. brucei* showcase unique biochemical mechanisms as efficient tools of adaptation that have been acquired during evolution as early divergent eukaryote. (**9**) One of the best examples of these tools is undoubtedly the biosynthesis and uptake of fatty acids (FAs). For a long time, trypanosomes were believed to rely exclusively on the host for the supply of the FAs required to produce phospholipids for their cellular membranes. (**9**) This has since been rebutted: trypanosomes also rely upon two different FAs biosynthetic pathways. (**9**) One consists of acyl carrier protein (ACP) and of a mitochondrial fatty acid synthase type II (FASII), of prokaryotic origin, which produces ∼10% of the overall FAs. (**10**) The other route is a highly specialised mechanism which relies on a unique plant-like alternative FA elongase-like ketoacyl synthase (ELO) located into the endoplasmic reticulum, which provides a substantial amount of myristate required for glycosylphosphatidylinositol (GPI)-anchor biosynthesis in the BSF. (**11,12**) These enzymes are finely regulated to synthesise specific FAs required, i.e., myristate for BSF and stearate for PCF, as the main building blocks to remodel the parasites’ lipid pool, hence membrane and morphology, during progression through the different life-cycle forms and host environments (i.e. pH, temperature, nutrients).(**13**)

Alongside with ELO and FASII, *T. brucei* also evolved a vast repertoire of desaturase and elongases for the *de novo* biosynthesis of unsaturated fatty acids (UFAs) and polyunsaturated fatty acids (PUFAs). (**14**) Most of UFAs and PUFAs are normally synthesised from the FAs and lipids that *T. brucei* take up from the host *via* a series of receptor- and/or endocytosis-mediated mechanisms. (**15–17**) Overall *T. brucei* are able to synthesise 75% of 16C and 18C

FAs in the cell, of which the 80% are unsaturated species. (**16**) Particularly, a large amount of linoleic acid (LA) (18:2) and docosapentaenoic acid (DPA) (22:5) and docosahexaenoic acid (DHA) (22:6), and lower levels of oleic acid (OA) (18:1) and 16C FAs have been found in BSF of *T. brucei*, compared to the host environment. (**16**) The synthesis of these PUFAs is possible through the combined activity of either an initial desaturase, i.e., Δ9-desaturase (**18**), or front-end desaturases, i.e., Δ6-, Δ5- and Δ4-desaturases (**11**), or methyl-end desaturases, i.e., Δ12- desaturase (**19**), as well as elongases (ELO1-4), that are required to work in a concerted and alternating manner to form both omega-6 (ω-6) and omega-3 (ω-3) PUFAs. (**20,21**) Some of these enzymes have been biochemically characterised (Tb927.7.4180, Tb927.7.4170, Tb927.7.4160, Tb927.5.4530, Tb927.2.3080, Tb927.8.6000) (**11,12,14,18–21**), however the way in which *T. brucei* uptake, synthesise and use very long chain PUFAs (VLC-PUFAs) remains not fully understood. Therefore, we explored the function and role of the putative Δ6- desaturase (Tb11.v5.0580) in both *T. brucei* PCF and BSF. Δ6-desaturases have been shown in other organisms, such as in fungi and algae, to be bifunctional enzymes able to insert a new double bond either on Δ^9,12^-18:2 or Δ^9,12,15^-18:3 to form Δ^6,9,12^-18:3 and Δ^6,9,12,15^-18:4, respectively. (**22**) Surprisingly, we discovered that this desaturase has instead a key role in the synthesis of 20C and 22C PUFAs, and that its activity is influenced by the type of environmental FAs available to the parasites. Moreover, we were able to show that the level of expression of this desaturase triggers the parasites’ lipid remodelling in response to environmental fat-sources availability. This finding gave insights on the plasticity of the FA pathway. *T. brucei* can use this biosynthetic machinery as a highly tuneable tool to re-shape the lipid content of the cell membrane and in turn their morphology. This is in response to the change in the host’s environment and nutritional availability, which reflects important metabolic adaptations between hosts. (**2**)

## Results

### *T. brucei* desaturase gene sequence identification and level of expression in wild type

In order to study the activity of the unexplored Tb-Δ6 desaturase in the kinetoplastid parasite *T. brucei,* the genomic Kinetoplastid database TriTrypDB was searched to select the sequence of the gene of interest. (**23**) In this process, two genes with almost identical sequences (99.8% identity) were found: Tb11.v5.0580, located on chromosome 11, and Tb927.10.7100, located on chromosome 10 (Figure S1A). To confirm whether we were looking at two real copies of the same gene located on two different chromosomes, or at a ghost copy of one single gene, a PCR was performed on the gDNA extracted from BSF *T. brucei* WT lister 427. No product (expected size 5651 bp) was obtained using specific primers targeting the region between Tb11.v5.0580 and the downstream gene (Tb11.v5.0579) on chromosome 11 (Figure S1B and Figure S1C). A PCR product of the expected size (977 bp) was instead obtained when using specific primers targeting the region between Tb927.10.7100 and the upstream gene (Tb927.10.7110) on the chromosome 10 (Figure S1D and Figure S1E). These results suggested that there is only one copy of the desaturase gene of interest, and that this gene is located on the chromosome 10. Furthermore, *via* qRT-PCR we were able to confirm that, as observed in previous RNAseq data set by Siegel *et al*. (**24**), the level of expression of *Tb-Δ6* gene is 5- fold higher (p = 0.0016) in PCF than in BSF WT (Figure S1F).

### Down- and up-regulation of Tb-Δ6 desaturase in procyclic and bloodstream forms of *T. brucei*

To investigate the essentiality and function of the putative Tb-Δ6 desaturase, overexpression (Δ6-OE) and knock-down (Δ6-KD) cell lines were generated in both *T. brucei* PCF and BSF. Tb-Δ6 expression was down-regulated *via* the tetracycline (Tet)-inducible RNA interference (RNAi) p2T7-177 vector (Δ6-KD), (**24**),(**25**) and upregulated *via* the Tet-inducible pLew100 vector (Δ6-OE). (**26**) Alongside these cell lines, an add-back (Δ6-OK) cell line was also generated in PCF, by transfecting Δ6-KD cells with the pLew100 vector. The stable integration of the transfected DNA in the genomic DNA was confirmed by PCR (Figure S2 and Figure S3). Furthermore, the overexpression, knock-down and add-back (only for PCF) of Tb-Δ6 with and without Tet-induction and compared with the WT controls were confirmed by qRT-PCR (Figure S3G and Figure S3H).

### Tb-Δ6 is associated with the mitochondrial compartment of *T. brucei* PCF and BSF

The cellular location of Tb-Δ6 was identified using the Tet-inducible overexpression vector pLew100, which express a C-terminal-6HA-tag on the Tb-Δ6 protein. Δ6-OE and WT cells for both PCF and BSF were harvested after 48 h of Tet-induction. Initial analysis by immunoblotting showed that Tb-Δ6 was expressed in both PCF (Figure S4A) and BSF (Figure S4B) and migrated according to the predicted size (∼49 kDa). The Tb-Δ6-HA protein was then detected by immunofluorescence microscopy using an anti-HA secondary antibody in the presence of Mito tracker and DAPI stains. The green, fluorescent punctate signals from Tb- Δ6-HA and the red fluorescence of the stained mitochondria showed an overlapping distribution in both BSF (Figure 1A, second row) and PCF (Figure 1B, second row) cells. This result suggested that Tb-Δ6 is mostly localised/associated with parts of the mitochondria.

**Figure 1.**
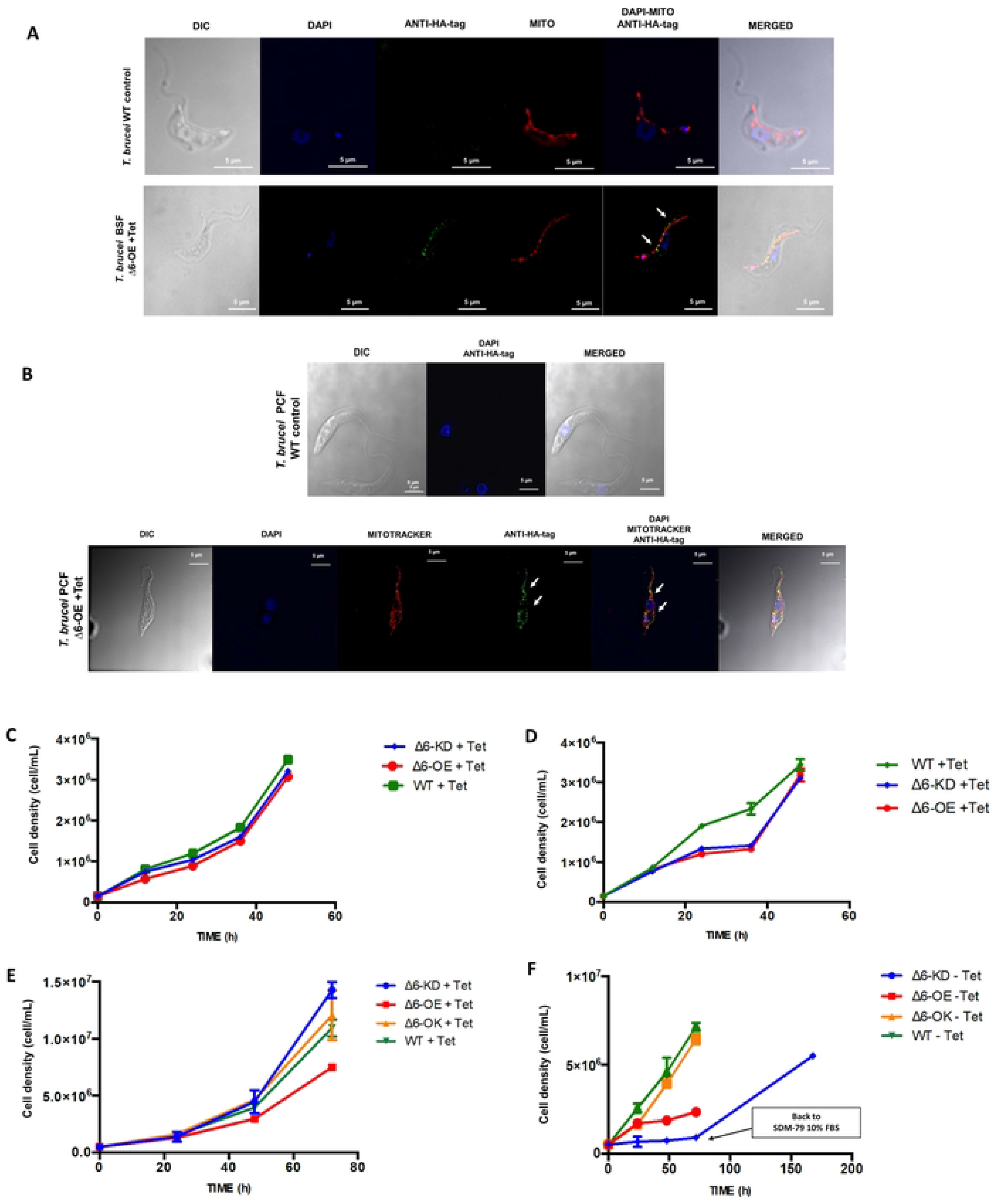
A, B) Tb-Δ6 cellular localisation in *T. brucei* BSF and PCF. Immunofluorescence microscopy images of *T. brucei* BSF WT control (**A**, first row) and PCF WT control (**B**, first row), and *T. brucei* BSF Δ6-OE (**A**, second row) and *T. brucei* PCF Δ6-OE (**B**, second row) fixed on poly-lysine coated slides stained with DAPI (blue signal), Mito tracker red (red signal), anti-HA tag (green signal) and imaged with DeltaVision Imaging System confocal microscope. The cells were grown for 48 h in HMI-11 with 10% FBS in the presence of tetracycline. The white arrows show that Δ6-desaturase (anti-HA tag green dots) is localised/associated with parts of the mitochondrial compartment (red, white arrows). Images were processed using softWoRx Explorer 1.3. **C, D) Growth curves of *T. brucei* BSF.** The graphs represent the growth curves over 48 h of *T. brucei* BSF WT control and *T. brucei* BSF Tb-Δ6 knock-down (Δ6-KD), and Tb-Δ6 overexpression (Δ6-OE) cells, when they are cultured in HMI-11 supplemented with 10% FBS (**C**) or with 5% FBS (**D**) in the presence of tetracycline as shown in the legend. **E, F) Growth curves of *T. brucei* PCF.** The graphs represent the growth curves over 72 h of *T. brucei* PCF WT control and *T. brucei* PCF Tb-Δ6 knock-down (Δ6-KD), Tb-Δ6 overexpression (Δ6-OE) and Tb-Δ6 add-back (Δ6-OK) cells, when they are cultured in SDM-79 supplemented with 10% FBS (**E**) or with 1.25% FBS (**F**) both in the presence of tetracycline, as shown in the legend. For all growth curves values are the mean of three independent biological replicates (n=3). Error bars represent the standard deviation of each mean. Statistical analysis was performed by GraphPad PRISM 6.0 using One-way ANOVA multiple comparisons based on a Tukey t-test with a 95% confidence interval.

### Knock-down and overexpression of Tb-Δ6 alter the growth rate depending upon the level of fatty acids available from Foetal Bovine Serum (FBS) in the media

Growth curves were performed to evaluate any significant effect of Tb-Δ6 genetic manipulation on *T. brucei* PCF and BSF cell proliferation. After 48 h in HMI-11 supplemented with 10% FBS, BSF WT, Δ6-KD and Δ6-OE cells showed no significant differences in their growth rate either in the presence (Figure 1C) or absence (Figure S5A) of Tet. The same was observed for PCF WT, Δ6-KD and Δ6-OE cells cultured for 72 h in SDM-79 supplemented with 10% FBS, either in the presence (Figure 1E) or absence (Figure S5C) of Tet. No significant differences were observed between the cell growth of WT control and Δ6-OK PCF under induction (Figure 1C), suggesting that the WT phenotype had been efficiently rescued from the initial knock-down of Tb-Δ6.

However, significant differences were observed when *T. brucei* PCF and BSF were cultured in low-fat media. Both the Δ6-OE and Δ6-KD BSF mutants revealed a slower cell growth over the first 24 h (̴ 2-fold slower than the WT) in the presence of Tet (Figure 1D). At this point, the growth reached a plateau, that was maintained for 12 h (Figure 1D). This lag and static growth phase were reproducibly observed numerous times. After a further 12 h, these cells returned to a growth rate similar to those of the WT.

The differences in the growth rate were even more evident in PCF. Δ6-OE PCF showed a slower growth rate (̴ 3-fold slower than WT), possibly due to a very low availability of suitable FAs from the reduced 1.25% FBS (Figure 1F). Δ6-KD PCF mutants showed a very slow and almost stationary growth (̴ 8-fold lower than WT, p < 0.0001) over the observed time period (Figure 1F). When Δ6-KD PCF cells were transferred back to SDM-79 with 10% FBS (Figure 1F), the cells were able to reach a final cell density similar to the one of Δ6-OK and WT control after 4 days (Figure 1F). No significant differences were observed for both BSF (Figure S5B) and PCF (Figure S5D) cell lines in absence of Tet in low-fat media.

### Effect of *T. brucei* Δ6-desaturase on the fatty acid content in *T. brucei* PCF and BSF

The results above showed that *T. brucei* proliferation changes according to Tb-Δ6 level of activity, developing upon the availability of extracellular fats in the media. To understand whether the FA metabolism was also altered in these conditions, the total FA content in PCF and BSF WT controls and mutants grown in high- and low-fat media, and in the presence or absence of Tet, was analysed *via* GC-MS.

#### The starting point: fatty acid profile of T. brucei PCF and BSF wild type

It is important to highlight that, when cultured in flasks, trypanosomes rely exclusively on the FBS contained within the media as the primary and only extracellular source of FAs, they are as free FAs, as well as neutral, phospholipids and sterol-esters. Analysis *via* GC-MS of the FAs present in the serum used in these experiments showed that stearic acid (18:0), oleic acid (18:1) and linoleic acid (18:2) represent 60% of the FAs present in the FBS, whereas only 13% is represented by PUFAs (**Error! Reference source not found.**). The remaining part is mostly represented by short and long chain saturated fatty acids (SAFAs) and by arachidonic acid (20:4) (**Error! Reference source not found.**). When reducing the level of FBS in the culture media, the cells will have a lower level of fats available for uptake and for *de novo* synthesis. To elucidate how the cells respond to genetic alteration and chemical manipulation of the media, the FA cellular content of both WT *T. brucei* PCF and BSF were analysed under standard culturing condition or rather in high-FBS (10%) media (Figures S6A and S7A). As expected, and largely in keeping with previous studies (**16**), myristic acid (14:0) and palmitic acid (16:0) were found to be present in both forms of *T. brucei*. Small amounts of 16:1 and 16:2, as well as 17:0, 17:1 and 17:2, were also detected (Figures S6A and S7A). Stearic acid (18:0), 18:2 and two species of 18:1 (Figures S6A and S7A) were the most abundant FAs, representing between 50-60% of the whole FA content in the cells (Figures S6A and S7A). Conversely, PUFAs were present at different ratios in BSF and PCF trypanosomes. It was possible to detect 20:4, 20:3 and 20:2, 22:6 (only detected in BSF cells), 22:5, 22:4 and 22:3 PUFAs, which accounted for 25-30% of the total FA content (Figures S6A and S7A) in both PCF and BSF. A lower amount of the same species (2:5, 22:4 and 22:3 PUFAs) were found in *T. brucei* PCF and BSF cultured in low-fat media (5% for BSF and 1.25% for PCF) (Figures S6B and S7B).

#### The fatty acid content of T. brucei PCF and BSF changes depending upon the level of Tb-Δ6-desaturase activity and the fatty acid sources in the media

The data shown above suggests that when BSF and PCF WT are exposed to a lower amount of FBS, despite no detrimental effects on the growth rate were detected, the FA content changes. To investigate this effect in conjunction with Tb-Δ6 knock-down and overexpression, the FA cellular content was analysed for Δ6-KD, Δ6-OE, Δ6-OK *T. brucei* PCF (Figure 2; Figures S8 and Figure S9) and Δ6-KD and Δ6-OE *T. brucei* BSF (Figure 2; Figures S10 and S11) grown in both high- and low-fat media in the presence and absence of Tet (Figures S12 and S13).

**Figure 2.**
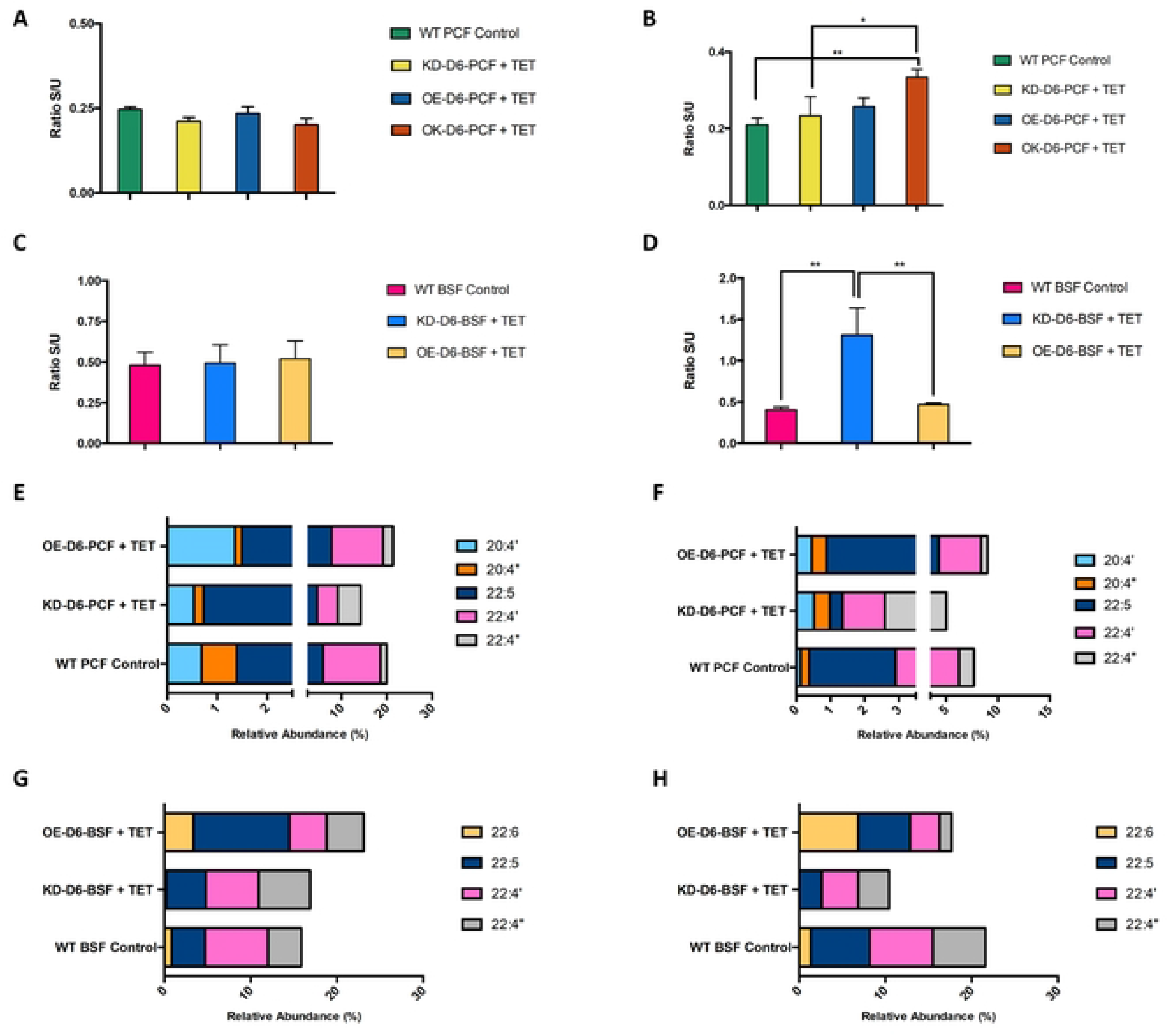
A, B, C, D) The ratio between saturated and unsaturated fatty acids in Tb-Δ6 genetically modified *T. brucei* PCF and BSF in high- and low-fat media. The bar charts show the ratio between the total amount of SAFAs (S) versus the total amount of the UFAs (U) produced by Tb-Δ6 genetically modified *T. brucei* PCF (KD-D6, OE-D6, OK-D6) and WT control in SDM-79 with 10% FBS (**A**) and in SDM-79 with 1.25% FBS (**B**), *T. brucei* BSF (KD-D6, OE-D6) and WT control in HMI-11 with 10% FBS (**C**) and in HMI-11 with 5% FBS (**D**) in the presence of Tet. Values are the mean of three independent biological replicates (n=3). Error bars represent the standard deviation of each mean. Statistical analysis was performed by GraphPad PRISM 6.0 using One-way ANOVA multiple comparisons based on a Tukey t-test with a 95% confidence interval, where ** is p ≤ 0.01 and * is p ≤ 0.05. **E, F, G, H) Comparison of 20C and 22C PUFAs between Tb-Δ6 genetically manipulated *T. brucei* PCF and BSF in high- and low-fat media.** The bar charts aim to give a visual summary of the difference in 20:4, 22:6, 22:5 and 22:4 PUFAs comparing the total relative abundance of each FA (X axis) between Tb-Δ6 genetically modified *T. brucei* PCF (KD-D6 and OE-D6) and WT control, when the cells are cultured in SDM-79 supplemented with 10 % (**E**) and 1.25% (**F**) FBS, and *T. brucei* BSF (KD-D6 and OE-D6) and WT control, when the cells are cultured in HMI-11 supplemented with 10 % (**G**) and 5% (**H**) FBS in the presence of tetracycline. All values for **E** and **F** are taken from **Figure S8** and **Figure S9**. All values for **G** and **H** are taken from **Figure S10** and **Figure S11**. <u>Note:</u> ‘ = first eluted isomer; “ = second eluted isomer.

Initially, the ratio between SAFAs (S) and UFAs (U) was calculated for *T. brucei* PCF (Figures 2A and 2B) and BSF WT controls (Figures 2C and 2D). No significant differences were observed between PCF WT and Δ6-KD, Δ6-OE and Δ6-OK PCF in high-fat media in the presence of Tet (Figure 2A). Surprisingly, the same outcome was obtained for WT and mutants cultured in low-fat media in the presence of Tet, except for Δ6-OK cells, which showed a higher ratio compared to the WT control (p = 0.0086) and Δ6-KD cells (p = 0.0297) (Figure 2B).

The same result was obtained for BSF WT control, Δ6-KD and Δ6-OE BSF cells grown in high-fat media in the presence of Tet (Figure 2C). Interestingly, a significant increase of SAFAs was detected for Δ6-KD BSF mutants in HMI-11 with 5% FBS compared to WT control (p = 0.0028) and to Δ6-OE BSF mutants (p = 0.0041), respectively (Figure 2D). On the other hand, Δ6-OE BSF cells showed an almost identical ratio to the one of the WT control (Figure 2D), and not different from the ratio in 10% FBS (Figure 2C).

To identify the classes of FAs that contributed to the observed change in the S/U, the FA cellular content was analysed in more detail. Interestingly, significant changes were observed in the relative abundance of 20:4, 22:4, 22:5 and 22:6 PUFAs, when comparing the WT control with the Δ6-KD, Δ6-OE and Δ6-OK *T. brucei* PCF and BSF cell lines, grown in either high- (10% FBS) or low- (1.25% or 5%) fat media in the presence of Tet.

The changes in these PUFAs detected in PCF showed the same trend in both high- (Figure 2E) and low-fat (Figure 2F) media. Particularly, in Δ6-KD mutants no significant differences were observed for the two isomers of 20:4 (Figure 2E; Figure S8B) and for 22:5 (Figure 2E; Figure S8C) compared to the WT control in high-fat media. On the other hand, one isomer of 22:4 decreased by 2.4-fold compared to the WT control (p < 0.0001) (Figure 2E, pink; Figure S8C), whereas the other isomer of 22:4 increased by 3.6-fold (p = 0.0055) when compared to the WT control (Figure 2E, grey; Figure S8C). Importantly, the second isomer of 22:4 was 2.3-fold higher (p = 0.0356) in Δ6-KD cells than in Δ6-OE cells in high-fat media (Figure 2E, grey; Figure S8C). In Δ6-OE mutants a biologically significant increase of 1.4-fold in 22:5 was observed compared to both WT control and Δ6-KD cells in high-fat media (Figure 2E; Figure S8C). Furthermore, 20:4 was 2-fold (p = 0.0431) and 2.5-fold (p = 0.0119) higher in Δ6-OE mutants than in the WT control and Δ6-KD mutants in high-fat media, respectively (Figure 2E, light blue; Figure S8B).

Some of these differences seemed to be more evident for PCF in low-fat media. One isomer of 20:4 was 4-fold (ns) higher in Δ6-KD mutants than in the WT control (Figure 2F, light blue; Figure S9B). Importantly, 22:5 was 6.8-fold (p = 0.0293) and 9.2-fold (p =0.0012) lower in Δ6-KD mutants than in the WT control and Δ6-OE cells, respectively (Figure 2F; Figure S9B). Furthermore, one isomer of 22:4 was 1.7-fold higher (p = 0.0300) in Δ6-KD mutants than in the WT control (Figure 2F, grey; Figure S9B). In low-fat media, Δ6-OE mutants showed a biologically significant increase of 22:5 compared to the WT (1.4-fold) (Figure 2F, Figure S9C), and a consequent decrease of the second isomer of 22:4 (2.4-fold) (Figure 2F, grey; Figure S9C).

As expected, Δ6-OK mutants did not show any significant difference compared to the WT control (Figure S8 and Figure S9), even in low-fat media where the ratio S/U was calculated to be higher than in the WT control (Figure 2B; Figures S8 and Figure S9).

Importantly, no production of 22:6 was detected in any of the PCF cell lines either in high- or low-fat media (Figure 2E and Figure 2F; Figures S8 and Figure S9). As expected, there were no significant differences between the WT control and the mutants in the absence of Tet (Figure S12).

In high-fat media, *T. brucei* BSF Δ6-OE was 22:5 2.9-fold (p = 0.0056; p=0.0122) and 2.4-fold higher than the WT control and Δ6-KD cells, respectively, in high-fat media (Fig 2G; Figure S10C). In contrast with PCF, 22:6 was detected in all BSF cell lines cultured under these conditions (Figures 2G and Figure 2H; Figure S10 and Figure S11). Particularly, 22:6 was 4.0- fold higher in Δ6-OE cells (p = 0.0006) and 4.0-fold lower in Δ6-KD than in the WT control in high-fat media (Figure 2G; Figure S10D).

When BSF cells were cultured in low-fat media, 22:4 and 22:5 were lower in both Δ6-KD and Δ6-OE mutants than in the WT control (Figure 2H; Figures S11C and Figure S11D). Particularly, in Δ6-KD mutants both isomers of 22:4 were 1.7-fold lower (p = 0.0026; p = 0.0004) than the WT control (Figure 2H; Figure S11D). Importantly, the second isomer of 22:4 was 2.5-fold higher in Δ6-KD mutants than in Δ6-OE cells (p = 0.0251) (Figure 2H, grey; Figures S11C and Figure S11D). On the other hand, 22:5 was 2.6-fold lower in Δ6-KD cells (p = 0.0046) compared to the WT (Figure 2H; Figure S11D).

Interestingly, 22:6 was not detected in Δ6-KD BSF mutants compared to the WT control and Δ6-OE mutants (p < 0.0001) (Figure 2H; Figure S11F). In the case of Δ6-OE mutants, one isomer of 22:4 was 4.3-fold lower in Δ6-OE mutants (p < 0.0001) compared to the WT control (Figure 2H, grey; Figure S11C), whereas the other isomer was 2-fold lower (p = 0.0004) than the WT control (Figure 2H, pink; Figure S11C). In addition, 22:5 showed a 1.4-fold decrease in Δ6-OE cells (biologically significant), when compared to the WT control (Figure 2H; Figure S11D). Importantly, a significant 5.0-fold increase in the amount of 22:6 was detected in Δ6- OE cells compared to the WT control (p = 0.0003) (Figure 2H; Figure S11F). No significant differences were identified in the absence of Tet, where the PUFA profile was very similar to the one of the cells induced with Tet (Figure S13A and Figure S13B). This is likely due to the effect of the minimal amount Tet often present in the FBS.

#### The ratio of product and substrate of Tb-Δ6-desaturase changes with its level of expression

As described above, the most consistent changes in the FA pool had been observed for the 20C and 22C PUFAs. This led to the hypothesis that 22:5 and/or 22:6 and 20:4 and/or 22:4 could be, respectively, the products (P) and substrates (S) of a multistep elongation-desaturation biotransformation, involving the Tb-Δ6 (Figure 3A). To test this hypothesis, the ratio (P/S) of products (22:5 and 22:6) (P) and substrate (20:4 and 22:4) (S) was determined for PCF and BSF in both high- and low-fat media (Figure 3B and Figure 3C respectively). A 1.4-fold increase and a 2-fold increase of P/S were detected in Δ6-OE PCF mutants compared to the PCF WT control (p = 0.0314, p = 0.0027) in high- and low-fat media, respectively, in the presence of Tet (Figure 3B and Figure 3C). A 3.2-fold decrease in P/S was detected for Δ6-KD PCF mutants compared to the WT PCF control (p = 0.0142) in low-fat media in the presence of Tet (Figure 3C). These results seem to suggest that to a higher level of products corresponds a higher level of expression of Tb-Δ6, whereas to a higher level of substrates corresponds a lower level of expression of Tb-Δ6. The observation was supported by the significant 6.2-fold increase in P/S detected in Δ6-OE cells compared to Δ6-KD cells (p = 0.0001) (Figure 3C).

**Figure 3.**
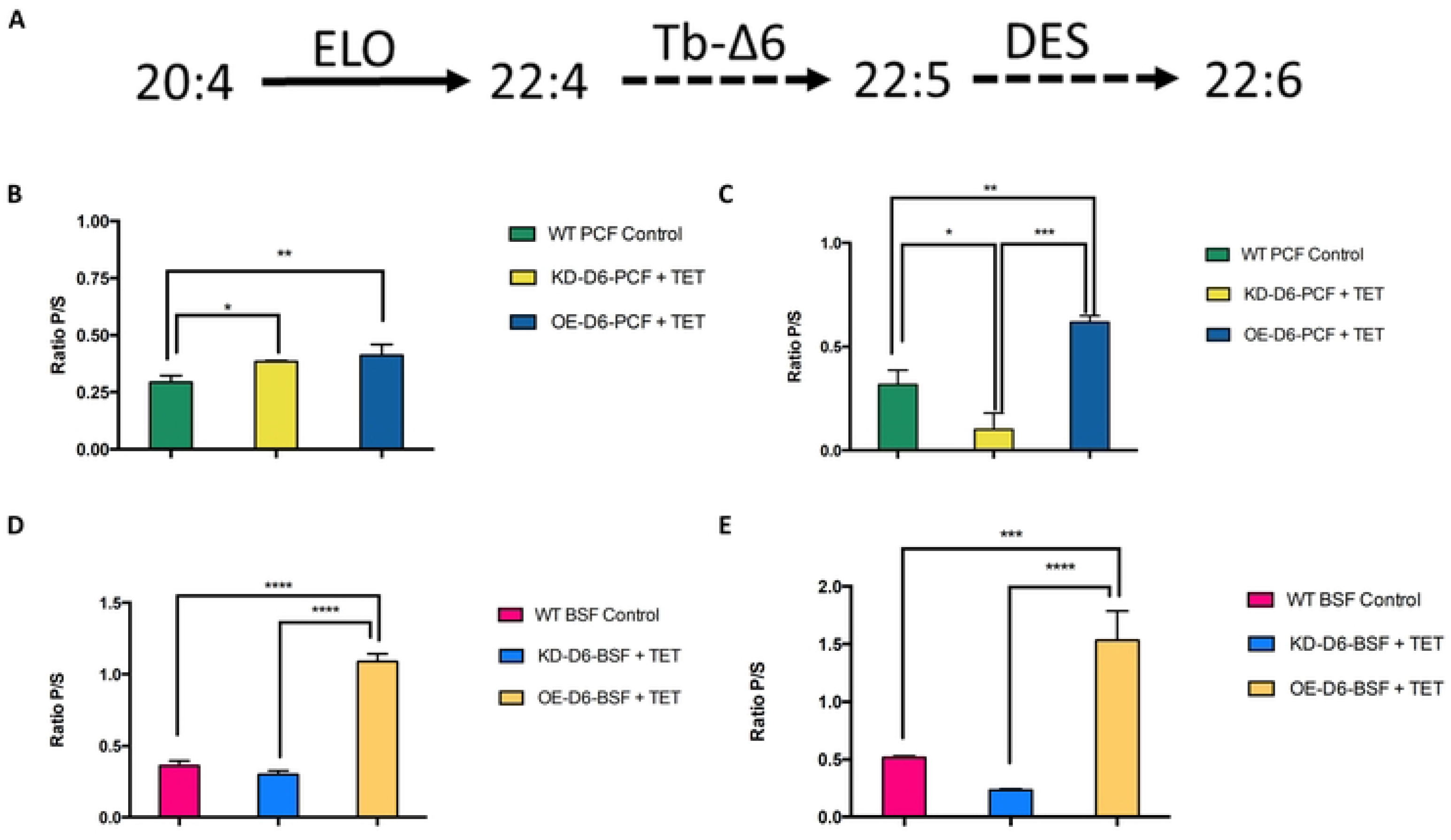
A) Hypothetical mechanism for the biotransformation of 20C and 22C PUFAs involving Tb-Δ6. The reaction scheme depicts one hypothetical mechanism for the formation of 22C PUFAs in *T. brucei*. In this case 20:4 is elongated to 22:4. The latter goes through desaturation by Tb-Δ6 to make 22:5, and finally 22:6 after further desaturation by DES. **B, C, D, E) The ratio between products (P) and substrates (S) of the biotransformation of 20C and 22C PUFAs in Tb-Δ6 genetically modified *T. brucei* PCF and BSF in high- and low-fat media.** The bar charts show the ratio between the total amount of hypothetical products (P) 22:5 and 22:6 PUFAs versus the total amount of the substrates (S) 20:4 and 22:4 PUFAs produced by Tb-Δ6 genetically modified *T. brucei* PCF (KD-D6 and OE-D6) and WT control in SDM-79 with 10% FBS (**B**) and with 1.25% FBS (**C**), and *T. brucei* BSF (KD-D6 and OE- D6) and WT control in HMI-11 with 10% FBS (**D**) and with 1.25% FBS (**E**). All values are the mean of three independent biological replicates (n=3). Error bars represent the standard deviation of each mean (±). All FAs were identified using GC-MS based upon retention time, fragmentation, and comparison with standards. Statistical analysis was performed by GraphPad PRISM 6.0 using 2-ways or One-way ANOVA multiple comparisons based on a Tukey t-test with a 95% confidence interval, where **** is p ≤ 0.0001, *** is p ≤ 0.001, ** is p ≤ 0.01 and * is p ≤ 0.05.

The same trend was observed for *T. brucei* BSF (Figure 3D and Figure 3E). A significant 3- fold increase of P/S was detected in Δ6-OE BSF mutants compared to the BSF WT control (p < 0.0001) in high-fat media and in the presence of Tet (Figure 3D). Under these conditions, a 3.7-fold change was detected in Δ6-OE BSF mutants compared to Δ6-KD BSF mutants (p < 0.0001) (Figure 3D). In the low-fat media, P/S decreased by 2.2-fold in Δ6-KD BSF cells compared to the WT BSF control (Figure 3E). Once again, this effect could be linked to the reduced activity of Tb-Δ6 upon scarce fat sources. Moreover, Δ6-OE BSF mutants showed a 3-fold and a 6.7-fold increase in the P/S ratio compared to the WT BSF control (p = 0.0003; p < 0.0001) and to Δ6-KD BSF mutants (p < 0.0001), respectively, under the same conditions (Figure 3E). Ultimately, these results confirmed that a greater level of products, i.e., 22:5 and 22:6, or substrates, i.e., 20:4 and 22:4, are linked to the overexpression and knock-down of Tb- Δ6, respectively.

### Supplementation of genetically modified *T. brucei* PCF and BSF with fatty acids confirms the substrate consumption and the product formation by Tb-Δ6-desaturase

To confirm that 22:5 and/or 22:6, and 20:4 and/or 22:4 are indeed the products and substrates respectively of Tb-Δ6, PCF and BSF WT and Δ6-KD and/or Δ6-OE cells were supplemented either with docosahexaenoic acid (Δ^4,7,10,13,16,19^-22:6) (DHA) or with arachidonic acid (Δ^5,8,11,14^- 20:4) (AA) in the presence and absence of Tet. Their growth rates and FA profiles under these conditions were subsequently analysed (Figure 4, Figure S17A and Figure S17B).

**Figure 4.**
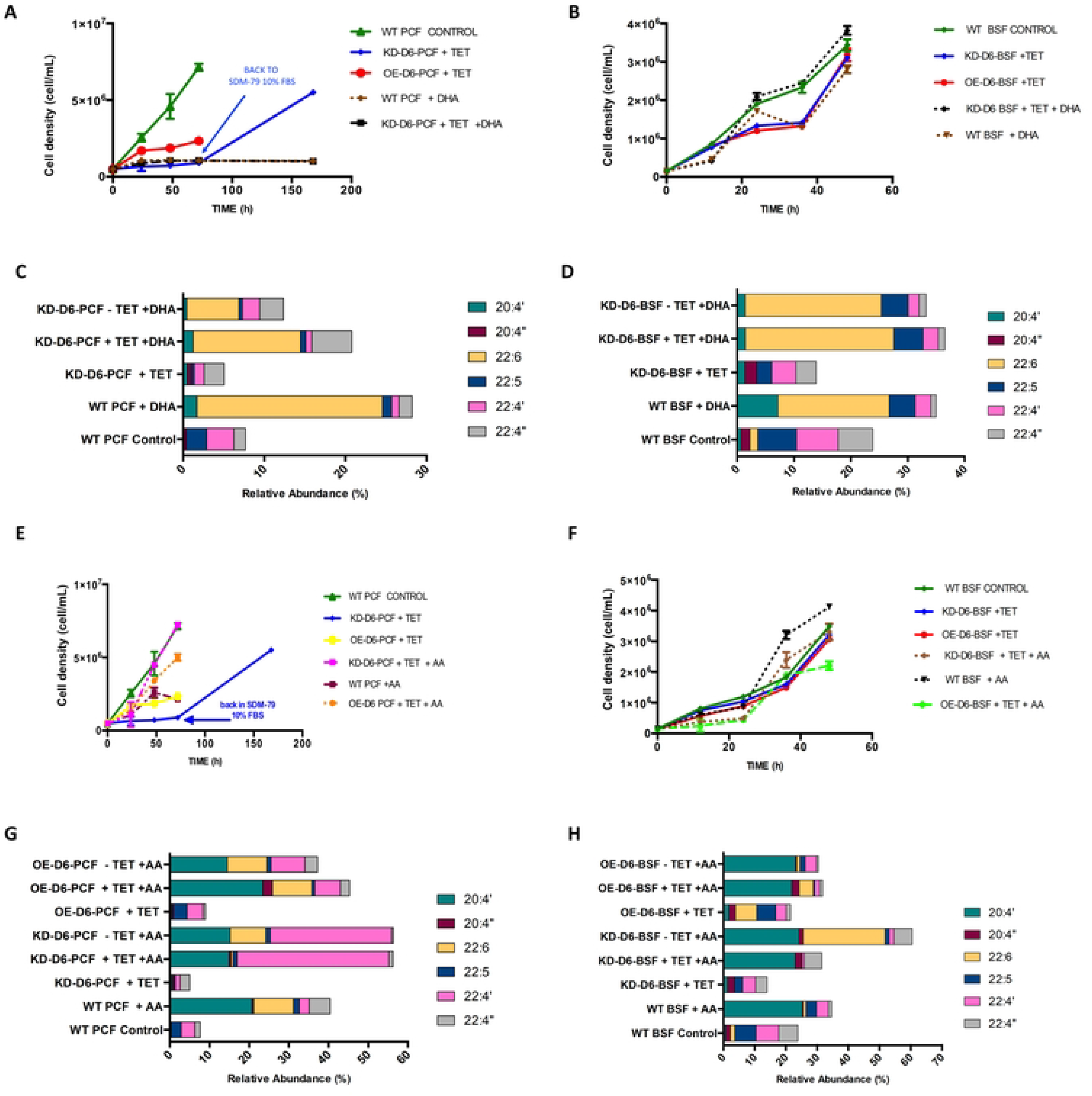
A, B) Growth curves of Tb-Δ6 genetically modified *T. brucei* PCF and BSF supplemented with DHA. The graphs represent the growth curves over 48 h of *T. brucei* PCF (**A**) and BSF (**B**) WT control and *T. brucei* PCF Δ6-desaturase knock down (KD-D6), when they are cultured in SDM-79 supplemented with 1.25% FBS (**A**) and HMI-11 supplemented with 5% FBS (**B**) (solid lines), and in SDM-79 supplemented with 1.25% FBS (**A**) and HMI-11(**B**) supplemented with 5% FBS added with 10 µM DHA (22:6) (dotted/dashed lines), in the presence of tetracycline as shown in the legend. **C, D) Comparison of 20C and 22C PUFAs between Tb-Δ6 genetically manipulated *T. brucei* PCF and BSF in low-fat media supplemented with DHA.** The bar charts aim to give a visual summary of the change in 20:4, 22:6; 22:5 and 22:4 PUFAs comparing the total relative abundance of each PUFA (X axis) between Tb-Δ6 genetically modified *T. brucei* PCF (**C**) and BSF **(D**) (KD-D6) and WT control, when the cells are cultured in SDM-79 supplemented with 1.25% FBS (**C**) and HMI-11 supplemented with 5% FBS (**D**) with or without 10 µM DHA and with or without tetracycline. All values are taken from **Error! Reference source not found.S14** and **Figure S15**. <u>Note:</u> ‘ = first eluted isomer; “ = second eluted isomer. **E, F) Growth curves of Tb-Δ6 genetically modified *T. brucei* PCF and BSF supplemented with DHA.** The graphs represent the growth curves over 48 h of *T. brucei* PCF (**A**) and BSF (**B**) WT control and *T. brucei* PCF Δ6-desaturase knock down (KD-D6) and overexpression (OE- D6), when they are cultured in SDM-79 supplemented with 1.25% FBS (**E**) and HMI-11 supplemented with 5% FBS (**F**) (solid lines), and in SDM-79 supplemented with 1.25% FBS (**E**) and HMI-11 (**F**) supplemented with 5% FBS added with 10 µM AA (20:4) (dotted/dashed lines), in the presence of tetracycline as shown in the legend. **G, H) Comparison of 20C and 22C PUFAs between Tb-Δ6 genetically manipulated *T. brucei* PCF and BSF in low-fat media supplemented with AA.** The bar charts aim to give a visual summary of the change in 20:4, 22:6; 22:5 and 22:4 PUFAs comparing the total relative abundance of each PUFA (X axis) between Tb-Δ6 genetically modified *T. brucei* PCF (**G**) and BSF **(H**) (KD-D6, OE-D6) and WT control, when the cells are cultured in SDM-79 supplemented with 1.25% FBS (**G**) and HMI-11 supplemented with 5% FBS (**H**) with or without 10 µM AA and with or without tetracycline. All values are taken from **Error! Reference source not found.S16** and **Figure S17**. <u>Note:</u> ‘ = first eluted isomer; “ = second eluted isomer. All values are the mean of three independent biological replicates (n=3). Error bars represent the standard deviation of each mean. Statistical analysis was performed by GraphPad PRISM 6.0 using 2-ways or One-way ANOVA multiple comparisons based on a Tukey t-test with a 95% confidence interval.

#### Growth rate of T. brucei PCF and BSF after supplementation with docosahexaenoic acid

After 72 h with DHA supplementation, both in the presence and absence of Tet, Δ6-KD PCF mutants and the PCF WT control showed an unexpected lag in growth (Figure 4A). The cells were not able to restore a normal growth rate even after 6 days of being cultured under these conditions. This was in contrast with what observed in the initial experiments, where Δ6-KD PCF cells restored their growth when transferred back into a fat-rich media after several days (Figure 1F). To explain the growth stall observed, and considering that DHA had not been detected in PCF trypanosomes so far, a cytotoxicity assay was also performed to exclude any toxic effect it may exert on PCF. No significant toxic effect was observed under the concentrations tested (Figure S14).

In contrast to PCF, Δ6-KD BSF supplemented with DHA rapidly reached a cell density equal to the one of the non-supplemented WT control, after dividing for 12 h at a slower rate than the WT control, than Δ6-KD and Δ6-OE mutants without DHA (Figure 4B). Δ6-KD mutants maintained this growth for a further 24 h. Moreover, BSF Δ6-KD mutants with DHA supplementation showed an increase in the cell density between the 12–36-hour time points compared to Δ6-OE cells grown in standard conditions (Figure 4B). After a 48-hour period of adaptation, all BSF cell lines were able to reach a very similar cell density (Figure 4B).

#### PUFA profile in T. brucei PCF and BSF after supplementation with docosahexaenoic acid

After 48 h with DHA supplementation, WT and Δ6-KD PCF cells internalised DHA at different levels (Figure 4C, Figure S15). The relative abundance of 22:6 in the WT cells was ∼23% (p < 0.0001), compared to the ∼13% in Δ6-KD cells in the presence of Tet (p < 0.0001) and 8% in the absence of Tet (p < 0.0001) (Figure 4C; Figure S15B). Under DHA supplementation, significant differences were observed for 20:4, 22:4 and 22:5 PUFAs (Figure 4C, Figure S15). Particularly, Δ6-KD PCF cells supplemented with DHA and induced with Tet showed a 10.1- fold (p < 0.0001) increase for one of the isomers of 20:4 (Figure 4C, pink; Figure S15C), and a 2.3-fold (biologically significant) increase in the second eluted isomer of 22:4 (Figure 4C, grey; Figure S15D), when compared to the WT control. In Δ6-KD cells, supplemented with DHA and induced with Tet, 22:5 was 3.9-fold (p = 0.0027) lower than in the WT control (Figure 4C; Figure S15E). On the other hand, when Δ6-KD cells supplemented with DHA were not induced, 22:5 was 6.4-fold (p = 0.0011) lower than the WT control (Figure 4C; Figure S15E).

WT and Δ6-KD BSF internalised DHA, which was detected at a relative abundance between ∼ 20-25% (Figure 4D, Figure S16B). Furthermore, in Δ6-KD BSF cells supplemented with DHA and both in the presence and absence of Tet 20:4 was 5.1-fold lower than in the WT cells supplemented with DHA (p = 0.0042; p = 0.0040) (Figure 4D, Figure S16C). The two species of 22:4 were respectively 2.8-fold and 5.2-fold (p < 0.0001) lower in Δ6-KD BSF cells, supplemented with DHA and induced with Tet, than in the WT control (Figure 4D, Figure S16D). On the other hand, 22:5 was ∼ 1.4-fold lower (biologically significant; p = 0.0305) in Δ6-KD BSF supplemented with DHA, both induced and non-, compared to WT control (Figure 4D, Figure S16E).

#### Growth rate of T. brucei PCF and BSF after supplementation with arachidonic acid

The growth rate of PCF trypanosomes in low-fat media supplemented with AA was observed over 72 h, in the presence and absence of Tet (Figure 4E, Figure S17C and Figure S17D). Upon Tet induction, Δ6-KD PCF mutants increased their growth by 8.1-fold over non-supplemented Δ6-KD mutants in low- and high-fat media, reaching similar cell density to the WT control (Figure 4E). In the presence of AA, the cell density of induced Δ6-KD PCF trypanosomes was 1.5-fold and 3.1-fold higher than induced Δ6-OE PCF cells supplemented and non-supplemented with AA, respectively (Figure 4E). On the other hand, the growth of induced and supplemented Δ6-OE PCF resulted in a 2.2-fold increase in cell density, but these cells did not grow as efficiently as the supplemented Δ6-KD trypanosomes (Figure 4E). Interestingly, the WT cells did not tolerate the excess of AA in the media: after 72 h the cell density was reduced by 3.4-fold compared to the WT control in the absence of AA (Figure 4E). In the absence of Tet, it appears that the supplementation with AA inhibits the growth of Δ6-OE cells (Figure S17C).

WT, Δ6-KD and Δ6-OE BSF trypanosomes supplemented with AA showed different cell density at each time point (Figure 4F). Once again, there were no significant differences in the growth rate among the cell lines cultured in the presence (Figure 4F), or absence of Tet (Figure S17D). AA supplementation had no effect on the growth of Δ6-KD and Δ6-OE BSF trypanosomes for the first 12 h (Figure 4F). After 24 h, a 2.4-fold decrease in cell density was detected for Δ6-OE and Δ6-KD BSF trypanosomes, when compared to the corresponding non-supplemented cells (Figure 4F). Δ6-KD BSF supplemented with AA reached the same cell density of the WT, Δ6-KD and Δ6-OE BSF trypanosomes grown in the absence of AA (Figure 4F and Figure S17D). On the other hand, AA supplementation reduced the final cell density of Δ6-OE BSF mutants by 1.6-fold at 48 h compared to Δ6-KD cells with AA (Figure 4F). Interestingly, WT trypanosomes supplemented with AA showed the highest cell density of all (Figure 4F). No differences were found between the cell lines either supplemented or not with AA, in the absence of Tet (Figure S17D).

#### PUFA profile in T. brucei PCF and BSF after supplementation with arachidonic acid

AA was efficiently internalised by the WT, Δ6-KD and Δ6-OE PCF cells, either in the presence or absence of Tet (Figure 4G, Figure S18A and Figure S18B). After supplementation with AA, it was possible to detect for the first-time the formation of 22:6 (p < 0.0001) in PCF (Figure 4G, Figure S18C). Interestingly, an 18.5-fold increase of 22:6 was detected in Δ6-KD cells supplemented with AA and in the absence of Tet compared to the induced Δ6-KD cells supplemented with AA (p < 0.0001) (Figure 4G, Figure S18C). Surprisingly, the same effect was not observed for induced and non-induced Δ6-OE mutants supplemented with AA, where instead the level of 22:6 remained very similar (̴10%) (Figure 4G, Figure S18C). Furthermore, one of the isomers of 22:4 was ̴14-fold higher in induced and non-induced Δ6-KD supplemented cells compared to the supplemented WT cells (p < 0.0001) (Figure 4G, Figure S18D). On the other hand, the second isomer of 22:4 was 4.8-fold and 11.2-fold lower in the induced and non-induced supplemented Δ6-KD cells than in the supplemented WT cells, respectively (ns) (Figure 4G, Figure S18D). In Δ6-OE mutants supplemented with AA, 22:4 was 2.6-fold higher in the presence of Tet and 3.4-fold higher in the absence of Tet than in the supplemented WT cells (p < 0.0001) (Figure 4G; Figure S18D).

BSF trypanosomes were also able to scavenge and internalise AA (Figure 4H, Figure S19A and S19B). One isomer of 22:4 was 5.2-fold (p = 0.0036; p = 0.0019) lower in Δ6-KD BSF cells, supplemented with AA, regardless of the induction with Tet, than in the supplemented WT cells (Figure 4H, Figure S19C). Moreover, the second species of 22:4 was 17-fold and 5.8-fold higher in induced and non-induced supplemented Δ6-KD cells than in Δ6- OE mutants grown in the same conditions (p = 0.0005; p = 0.0029 p = 0.0015; p = 0.0003) (Figure 4H, Figure S19C). Supplemented and induced Δ6-OE BSF cells revealed a 6-fold (p = 0.0001) decrease in 22:4, when compared to the non-supplemented WT control (Figure 4H, Figure S19C).

22:5 largely decreased when Δ6-KD cells, in the presence of AA and Tet, were compared to WT control (p = 0.0008) (Figure 4H, Figure S19D). Moreover, 22:5 was 14.2-fold lower (p < 0.0016) in induced and supplemented Δ6-OE mutants than in the non-supplemented WT, whereas when Δ6-OE mutants were not induced, 22:5 was 3.8-fold lower (p = 0.0096) than the WT control (Figure 4H, Figure S19D). Interestingly, it was possible to see production of 22:6 also in Δ6-KD cells upon supplementation with AA and without Tet induction (p < 0.0001) (Figure 4H, Figure S19E). No 22:6 was produced by the induced and supplemented Δ6-KD cells (p < 0.0001). Moreover, the level of 22:6 was 7.4-fold higher in the supplemented and induced Δ6-OE trypanosomes than in the supplemented WT cells (Figure 4H; Figure S19E). Furthermore, Δ6-OE cells, supplemented with AA and in presence of Tet, contained 3.4-fold more 22:6 than the same cell line in absence of Tet (p = 0.0353) (Figure 4H, Figure S19E).

It is important to highlight that both under DHA and AA supplementation PCF and BSF WT cells showed lower level of 18:1 (0.0001<p<0.0017; 0.0079<p<0.0161) and 18:2 (0.0099<p<0.0259;p = 0.0010) while presenting higher level of 18:0 (p = 0.0135; p = 0.0078) compared to PCF and BSF WT cells cultured in 10% FBS (Figure S20). This is a clear mechanism adopted by the cells to compensate for the high amount of PUFAs, not normally present in FBS (Table S1), but available under PUFA supplementation, when Tb-Δ6 is expressed at normal level.

### Determining the double bond position in the fatty acid product 22:6 formed in PCF after supplementation with AA

The results described above seemed to strongly support the hypothesis that 22:6 may be the final product of a series of two desaturation (one of which involves Tb-Δ6) and elongation reactions in *T. brucei*. This was particularly evident in *T. brucei* Δ6-KD PCF and BSF supplemented with AA. To define the exact position/geometry of the double bond inserted on the acyl-chain by Tb-Δ6, dimethyl disulfide (DMDS) derivatisation of the double bond was performed to obtain the DMDS-22:6 derivative (see Materials and Methods). The DMDS reaction was carried out on FAME mixtures from Δ6-KD PCF supplemented with AA, with and without Tet, to allow thorough analysis of the final products by GC-MS. The very complex mixture of products obtained at the end of the DMDS reaction made the assignment of the double bond position challenging. (**27**) The DMDS derivative of a DHA (22:6) standard showed a broad peak eluting at 36 min with an intact mass of 436 m/z as expected. The mass spectra revealed a specific fragmentation pattern, including diagnostic 147 m/z, 189 m/z and 89 m/z fragments that correspond respectively to DMDS addition at C4, C7 and C19 of the alkyl chain (**Error! Reference source not found.**A).(**27**) These same fragments were found in the cell extract samples. It was possible to identify a broad peak eluting at 35.7 min. This was a mixture of three different DMDS-PUFAs present in the sample. The characteristic fragments of 147 m/z and 288 m/z were assigned to Δ^4,7,10,13,16,19^-22:6, being respectively the addition of DMDS on C4 and C5 (Figure S21B, purple). The specific fragments of 189 m/z and 131 m/z were assigned to Δ^7,10,13,16^-22:4, being respectively the addition of DMDS on C16 and C7 (Figure S21B, orange).

### Genetic manipulation of Tb-Δ6-desaturase causes specific lipid modifications in *T. brucei* PCF and BSF in low-fat media

The differences in the PUFA profile revealed so far led us to investigate the possible phospholipid changes in the various differentially expressing Tb-Δ6 cell lines cultured under various FA sources in the media.

Lipidomic studies were carried out by electrospray ionization tandem mass spectrometry (ESI-MS/MS) to identify the differences in the phospholipid species between WT control cells and Δ6-KD and Δ6-OE mutants in PCF and BSF in both high- and low-fat media in the presence of Tet (Figure 5; Figure 6; Figure S22-S25). The lipids species were identified by comparison with the ones listed in the extensive work done by Richmond et al.,(**28**) and with structures, exact masses and fragmentation profiles of the various molecules collected in LIPID MAPS® database.

**Figure 5.**
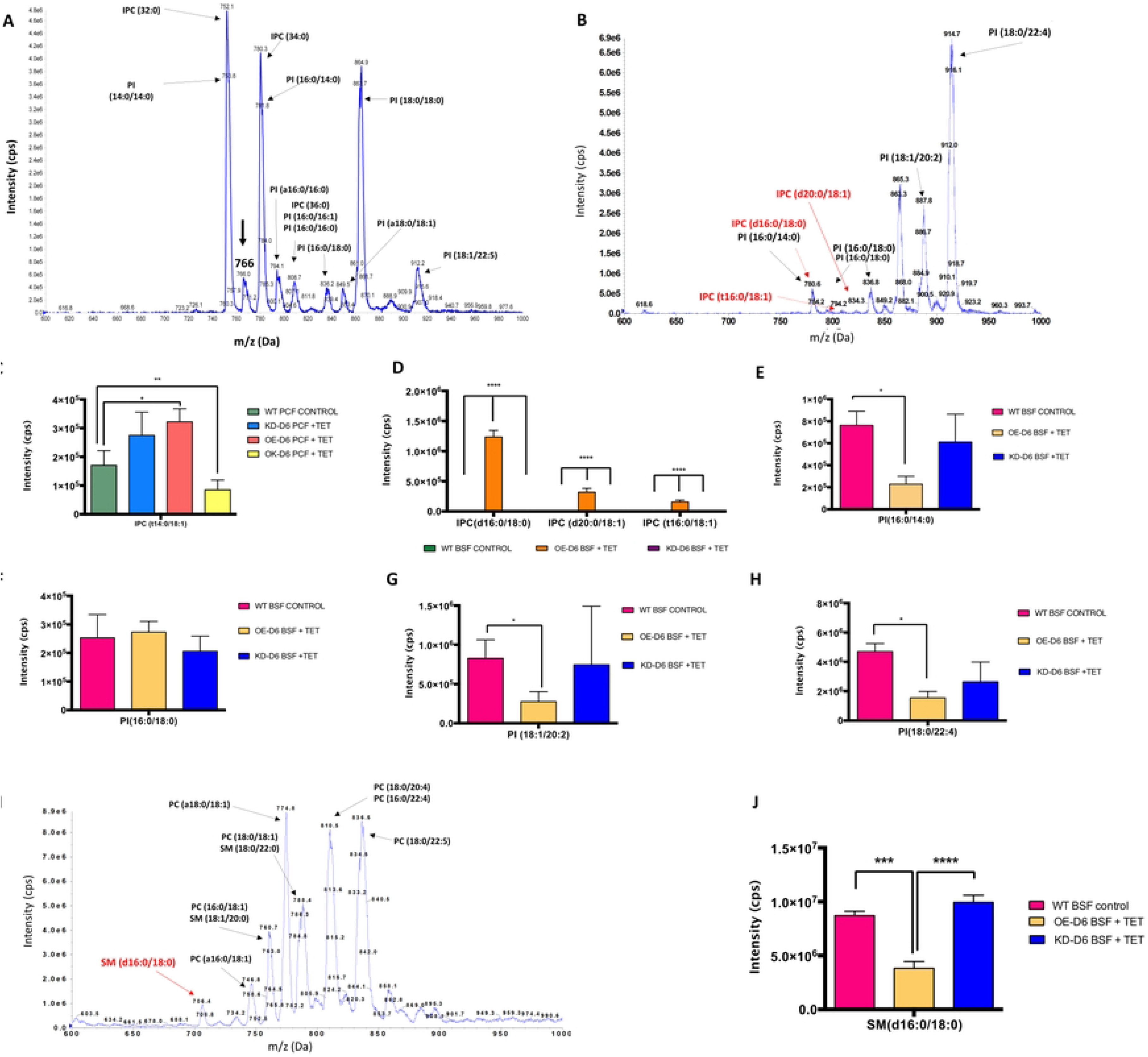
ESI-MS/MS of PI-containing lipids for genetically manipulated *T. brucei* PCF (A) and BSF (B) overexpressing Tb-Δ6 and grown in low-fat media. A) Shows representative spectra of PI- containing lipids obtained by scanning for parent ion of 241 m/z for Δ6-KD PCF grown in SDM-79 with 1.25% FBS in the presence of Tet. The species of interest is labelled (black) and highlighted by an arrow (black). The major PI and IPC species are also annotated.(**28**) a, alkylacyl. **B)** The spectrum shows PI-containing lipids and IPCs obtained by scanning for parent ion of 241 m/z for Δ6-OE BSF grown in HMI-11 with 5% FBS in the presence of Tet. The species of interest are labelled (PIs black, IPCs red) as reported in the text and highlighted by arrows (PIs black, IPCs red). d, sphingosine; t, 1,2,3-trihydroxy sphingosine. C-H) ESI-MS/MS quantification of PI-containing lipids for genetically manipulated *T. brucei* PCF (C) and BSF (D-H) overexpressing Tb-Δ6 and grown in low-fat media. The bar chart shows the difference in IPC (**C** and **D**) and PI (**E-H**) species (X axis) and the normalised intensity (Y axis, cps) found in Tb-Δ6 genetically modified *T. brucei* PCF (**C**) and BSF (**D-H**) OE-D6, KD-D6 and WT control, when the cells are cultured in SDM-79 supplemented with 1.25% FBS (**C**) and HMI-11 supplemented with 5% FBS (**D-H**), in the presence of tetracycline as shown in the legend. The relative intensities of PIs and IPCs were normalised against the intensity of PI (15:0/18:1(d7)) at 847.13 m/z contained in SPLASH internal standard. I) ESI-MS/MS spectra of PC-containing lipids for Tb-Δ6 genetically manipulated *T. brucei* BSF overexpressing Tb-Δ6 and grown in low-fat media. The spectrum shows PC-containing lipids obtained by scanning for parent ion of 184 m/z for WT control BSF grown in HMI-11 with 5% FBS. The SM of interest is labelled as reported in the text and highlighted by an arrow (red). The major SM and PC species are also annotated.^124^ a, alkylacyl. J) ESI-MS/MS quantification of PC-containing lipids for Tb-Δ6 genetically manipulated *T. brucei* BSF overexpressing Tb-Δ6 and grown in low-fat media. The bar chart shows the difference in SM species at 706 m/z (X axis) and the normalised intensity (Y axis, cps) found in Tb-Δ6 genetically modified *T. brucei* BSF (KD-D6 and OE-D6) and WT control, when the cells are cultured in HMI-11 supplemented with 5% FBS, in the presence of tetracycline as shown in the legend. The relative intensities of SM were normalised against the relative intensities of SM (d18:1/18:1(d9)) at 738.12 m/z contained in SPLASH internal standard. Values from the all bar chats are the mean of three independent biological replicates (n=3). Standard deviation of each mean (±) is calculated for the normalised intensities. Statistical analysis was performed by GraphPad PRISM 6.0 using One-way ANOVA multiple comparisons based on a Tukey t-test with a 95% confidence interval, where **** is p ≤ 0.0001, *** is p ≤ 0.001, ** is p ≤ 0.01 and * is p ≤ 0.05.

**Figure 6.**
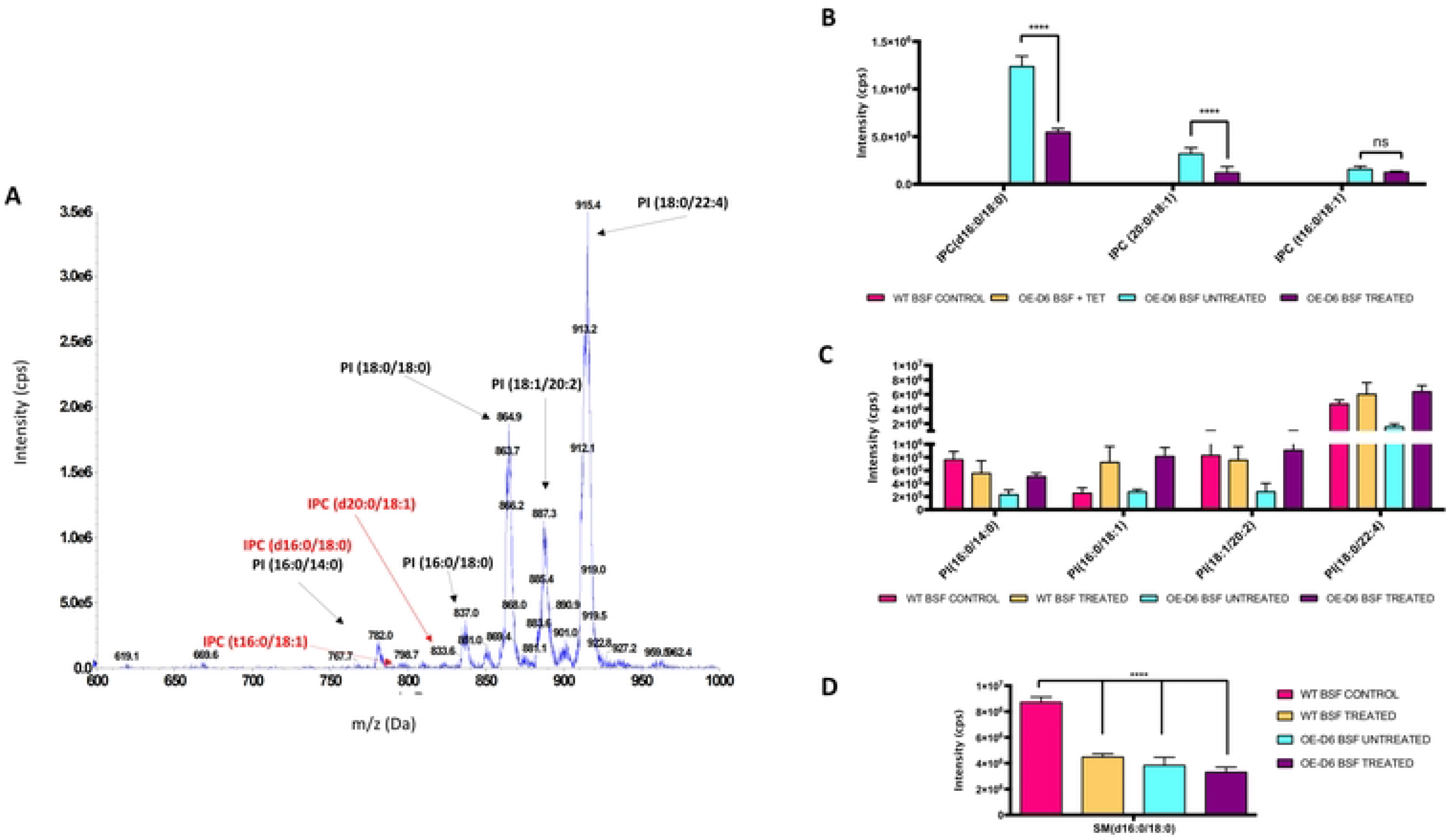
A) ESI-MS/MS of PI-containing lipids for Tb-Δ6 genetically manipulated *T. brucei* BSF grown in low-fat media treated with EC_10_ of clemastine fumarate. The spectrum shows PI-containing lipids by scanning for parent ion of 241 m/z for Δ6-OE BSF grown in HMI-11 with 5% FBS in the presence of Tet and treated with EC_10_ of clemastine fumarate. The IPC species of interested are labelled in red and highlighted with a red arrow. The PI species are annotated in black and pointed by black an arrow.(28) B-D) ESI-MS/MS quantification of IPCs, PIs and SM in Tb-Δ6 genetically manipulated *T. brucei* BSF in low-fat media after treatment with EC_10_ of clemastine fumarate. The bar chart shows the difference in IPC (**B**), PI (**C**) and SM (**D**) species (X axis) and the normalised intensity (Y axis, cps) found in Tb-Δ6 genetically modified *T. brucei* BSF OE-D6 and WT control, when the cells are cultured in HMI-11 supplemented with 5% FBS, in the presence of tetracycline, and treated or not with EC_10_ (0.61 µM calculated with GraphPad from the EC_50_) of clemastine fumarate, as shown in the legend. The relative intensities of PIs and IPCs were normalised against the intensity of PI (15:0/18:1(d7)) at 847.13 m/z contained in SPLASH internal standard. The relative intensities of SM were normalised against the relative intensities of SM (d18:1/18:1(d9)) at 738.12 m/z contained in SPLASH internal standard. Values are the mean of three independent biological replicates (n=3). Standard deviation of each mean (±) is calculated for the normalised intensities. Statistical analysis was performed by PRISM 6 using 2-way ANOVA multiple comparisons based on a Tukey t-test with a 95% confidence interval.

#### Overexpression of Tb-Δ6-desaturase causes production of inositolphosphorylceramide in T. brucei BSF upon low-fat media

ESI-MS/MS analysis of the lipid extracts obtained from the various PCF and BSF cell lines induced with Tet revealed no significant differences in the lipid profiles in positive and negative survey scans for BSF and PCF WT control grown in high- and low-fat media. However, specific changes in inositolphosphorylceramide (IPC) were detected in PCF mutants in low-fat media (Figure 5A), alongside with the unexpected formation of IPC in Δ6-OE BSF in low-fat media (Figure 5B). This finding was surprising, provided that IPC biosynthesis has only been observed in the stumpy and procyclic forms of *T. brucei.*(**29**), (**30**)

Starting with PCF, the scans for the parent ion of 241 m/z revealed a change in intensity of the peak at 766 m/z (Figure 5A). In this case, an increase of 1.6-fold (biologically significant) and of 1.8-fold (p = 0.0405) were detected in Δ6-KD and Δ6-OE cells compared to the WT control, respectively (Figure 5C). To assign the 766 m/z peak to a specific IPC, ESI-MS/MS daughter ion fragmentation was performed in the range 120-800 m/z for each sample, identifying the 766 m/z peak as IPC (t14:0/18:1) (where t- indicates the 1,3,4-trihydroxy 14:0 chain) (Figure S22A). On the other hand, no significant differences were observed for IPC (t14:0/18:1) in high-fat media (Figure S22B), and for the other major IPC species present in PCF in low-fat media (Figure S22C-S22F).

For BSF *T. brucei*, scans for the parent ion of 241 m/z revealed an increase of the intensity of some peaks in the range 780-840 m/z only in the induced Δ6-OE BSF cells in low-fat media (Figure 5B) compared to the WT control (Figure S23A) and Δ6-KD cells (Figure S23B). No production of IPC was detected in any of the BSF cell lines in high-fat media (Figure S23C- S23E). In low fat-media, IPC (d16:0/18:0) and PI (16:0/14:0) at 780 m/z (Figure 5B, Figure 5D and Figure 5E; Figure S24A), IPC (t16:0/18:1) at 794 m/z (Figure 5B-5D; Figure S24B), IPC (d20:0/18:1) at 834 m/z and PI (16:0/18:0) at 837 m/z (Figure 5B, Figure 5D and Figure 5F; Figure S24C) were identified. Moreover, a difference in the intensity of the peaks was found for PI (18:1/20:2) at 887 m/z (Figure 5B and Figure 5G) and PI (18:0/22:4) at 914 m/z (Figure 5B-5H), previously identified by Richmond et al. (**28**) As IPC phospholipids contain a ceramide anchor, which is also present in the choline-containing sphingolipid SM, normally produced in BSF (**29**),(**30**), the idea was that, if Δ6-OE mutants were producing IPC under Tb- Δ6 overexpression and reduced fat media, a reduction of SM biosynthesis and an alteration in some PI species might be observed due to the redirection of the ceramide anchor and use of the PI head group to facilitate IPC synthesis. Hence, precursors ion scans for choline-containing phospholipids at 184 m/z were performed (Figure 5I; Figure S25A and Figure S25B). Indeed, a decrease of 2.3-fold (p = 0.0001) and of 2.6-fold (p < 0.0001) of SM (d16:0/18:0) at 706 m/z was observed in Δ6-OE mutants in the presence of low-fat media compared to the WT control and induced Δ6-KD cells, respectively (Figure 5I and Figure 5J; Figure S25A and Figure S25B). As expected, a decrease in some of the PI species was also detected: PI (16:0/14:0), PI (18:1/20:2) and PI(18:0/22:4) were 3.0-fold (p = 0.0190; ns; p = 0.0105) lower in Δ6-OE mutants in the presence of low-fat media compared to the WT control (Figure 5E-5H; Figure S25A and Figure S25B). This phenomenon was not observed in BSF mutants and control grown in high-fat media, because of the absence of IPC, as expected (Figure S25C and Figure S25F).

#### Clemastine fumarate reduces IPC production in T. brucei BSF overexpressing Δ6-desaturase in reduced fat media

To assess whether the IPC production by Δ6-OE BSF mutants in low-fat media could be reduced/inhibited, we used the leishmanial IPC synthase inhibitor clemastine fumarate.(**31**) Firstly, clemastine fumarate was tested in a dose-response assay against *T. brucei* BSF WT, used as a control, and Δ6-OE BSF mutants induced for 48 h with Tet, in low-fat media. It was found that clemastine fumarate was active against *T. brucei* BSF WT with an EC_50_ 6.76 ± 0.15 µM (Figure S26A) and against induced Δ6-OE BSF trypanosomes with an EC_50_ 5.45 ± 0.30 µM (Figure S26B). The non-induced Δ6-OE BSF were also tested. They gave a value of EC_50_ 6.89 ± 0.26 µM (Figure S26C). A non-lethal concentration of clemastine fumarate (EC_10_ 0.61 µM) was then used to treat WT and Tet induced Δ6-OE BSF trypanosomes. The level of IPC was found to be diminished relative to PI compared to the corresponding untreated cell line (Figure 6A). Particularly, IPC (d16:0/18:0) was reduced by ∼2.3-fold (p < 0.0001) and IPC (d20:0/18:1) by ∼2.7-fold (p < 0.0001), whereas IPC (t16:0/18:1) was only diminished by 1.3- fold (ns) (Figure 6B). Interestingly, some of the PI species in Δ6-OE cells treated with clemastine fumarate, were restored to almost the same level as either the treated or untreated WT cells (Figure 6C). Thus, one may expect that clemastine fumarate inhibition of IPC formation would have allowed the ceramide to be re-utilised for SM synthesis. Unexpectedly, this was not the case, as the SM (d16:0/18:0) was ∼2-fold (p < 0.0001) lower in both treated and untreated induced Δ6-OE cells, and it was still 2-fold (p < 0.0001) lower in treated WT cells compared to the WT untreated control (Figure 6D).

#### The effect of the overexpression of Tb-Δ6 and of the formation of IPC on the cell morphology of T. brucei BSF

The results reported above imply that the genetic manipulation of Tb-Δ6, and particularly the overexpression of Tb-Δ6, under reduced fat conditions, affect the lipid composition of the membranes resulting in the formation of IPC. This is in accordance with the data showing that PCF *T. brucei* express higher level of Tb-Δ6 gene compared to BSF *T. brucei*. (**24**) It is also important to highlight that the *T. brucei* stumpy form, which overexpress Tb-Δ6, are indeed able to make IPC. (**29**) This might suggest that Tb-Δ6 and IPC, are inter-connected by a cascade of metabolic events, which reflects upon lipid remodelling, and ultimately on an alteration of the cells at the morphological level. We were in fact able to observe that when BSF *T. brucei* are overexpressing Tb-Δ6 and synthesising IPC in low fat media, the overall cell morphology is not elongated, as it would be expected for the slender form, but started assuming the appearance of a distinctly “plumpy” structure at 72 hours, similar to stumpy morphology (Figure 7). (**29**) Additionally their cell growth (Fig 1F), shows that there is hardly any cell-division. This implies that differential expression of both Tb-Δ6 and IPC synthase collectively might participate to a cascade of events that ultimately causes the morphological and biochemically transition from one life-stage to the other. To Test this hypothesis, immunofluorescence microscopy detection of the protein associated with differentiation 1 (PAD1), a molecular transducers of differentiation signals highly expressed in the stumpy forms, was performed by using anti-α-PAD1 primary antibody. (**32**) Expectedly, the stumpy *T. brucei* parasites, used as a positive control, showed their typical shorten and flattened shape as well as an intense signal for PAD1 on the cell surface (Figure 7, first row). (**32**) In keeping with studies by Matthews *et al*., WT negative control slender form of trypanosomes showed low signal for PAD1 (Figure 7, second row). (**29,32**) The same was observed for Tet induced Δ6-OE mutants grown in HMI-11 with 10% FBS (Figure 7, third row). Contrarily, Δ6-OE cells grown in low-fat media under Tet induction for 48 h, showed the presence of PAD1 on the cell surface and, particularly, in some peripheral areas (Figure 7, fourth row). After 72 h the PAD1 signal was found to be more intense and more broadly distributed on the cell surface (Figure 7, fifth row), as well as a more evident enlarged/flattened cell morphology and shortened flagellum.

**Figure 7.**
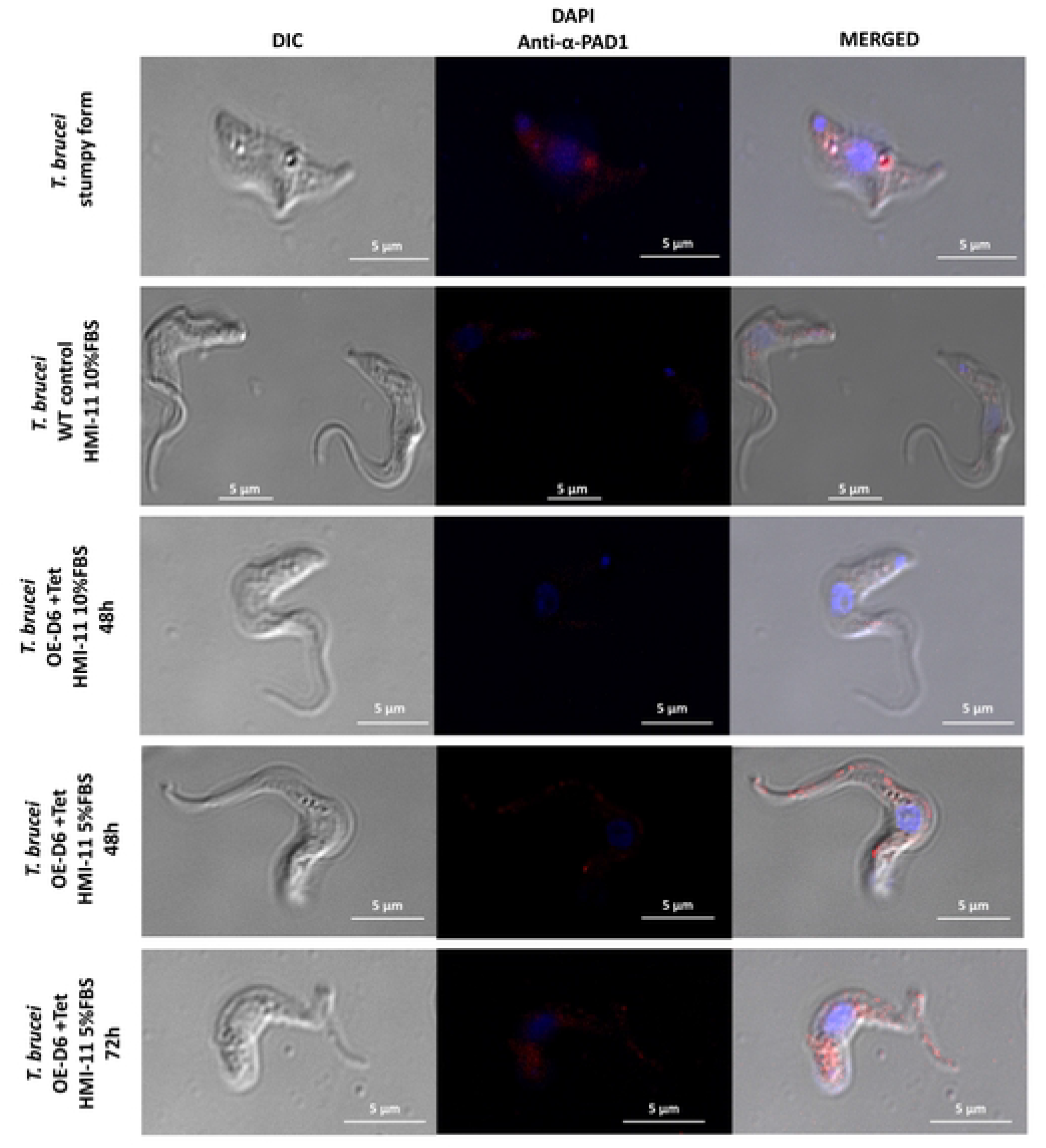
*T. brucei* BSF cell morphology changes when the cells overexpress Tb-Δ6 in low-fat and produce IPC. Immunofluorescence microscopy images of *T. brucei* BSF stumpy form (positive control) (first row), WT slender form (second row) and *T. brucei* BSF Δ6-OE (third, fourth and fifth rows) fixed on poly-lysine coated slides, stained with DAPI (blue signal) and anti-α-PAD1 antibody (red signal, **B**) and imaged with DeltaVision Imaging System microscope. The controls cells were grown for 48 h in HMI-11 with 5% FBS. Δ6-OE were grown in the same conditions in the presence of tetracycline for 48 h (fourth row, **B**) or 72 h (fifth row, **B**). Images were processed using softWoRx Explorer 1.3.

## Discussion

In this study, the essentiality of Tb-Δ6 in the biotransformation of PUFAs was elucidated, by genetic manipulation in *T. brucei* PCF and BSF, and by complementation of the level of essential fat sources in the media. Phenotypical responses were revealed in manipulated *T. brucei* after alteration of the expression level of Tb-Δ6, upon high and low level of appropriate FAs in the media. (**33,34**) These were, in fact, the first signals of significant metabolic changes within the cells. Modulation of the growth rate, together with visible morphological changes to the overall shape of the cells were detected as consequence of a process of adjustment of the biosynthesis of FAs and lipids, hence of remodelling of the lipid pool, as finely regulated tools used by the cells to adapt to the changes in the cellular environment. These mechanisms showed to be part of the adaption strategy implemented by the parasites to respond to the alteration of the activity of this essential desaturase, which was found to be fundamental for the normal cellular growth, by the reliance upon the level of appropriate fat sources, *i.e*., FBS, AA, DHA. In fact we demonstrated that the ability of trypanosomes to finely regulate the uptake and metabolism of FAs to achieve an appropriate redistribution of SAFAs and UFAs, upon genetic manipulation of Tb-Δ6 and upon the availability of the FA sources affects the biosynthetic pathway of PUFAs directly. (**35,36**) Significant differences in the 20C and 22C PUFAs were detected, by ultimately allowing the true activity of Tb-Δ6 to be unveiled. We found that this desaturase has a key role in the synthesis of essential ω-3 VLC-PUFA, namely 22:5 and 22:6, by working as Δ4-ω-end acyl-CoA desaturase, and in concerted action with other endogenous desaturases and elongases (Figure 8A). Furthermore, the HA-tagged Tb-ω-end-Δ4 (or Tb-Δ6) was detected to be localised in the mitochondrial compartment. The patchy signal obtained may suggest that the desaturase is localised in the mitochondrial contact sites with the ER. (**37**) This would clarify how the desaturated product of Tb-ω-end-Δ4 (or Tb-Δ6) from the mitochondria would get in contact with the elongase (ELO 4) present in the ER to be elongated to the very final product (Figure 8A).

**Figure 8.**
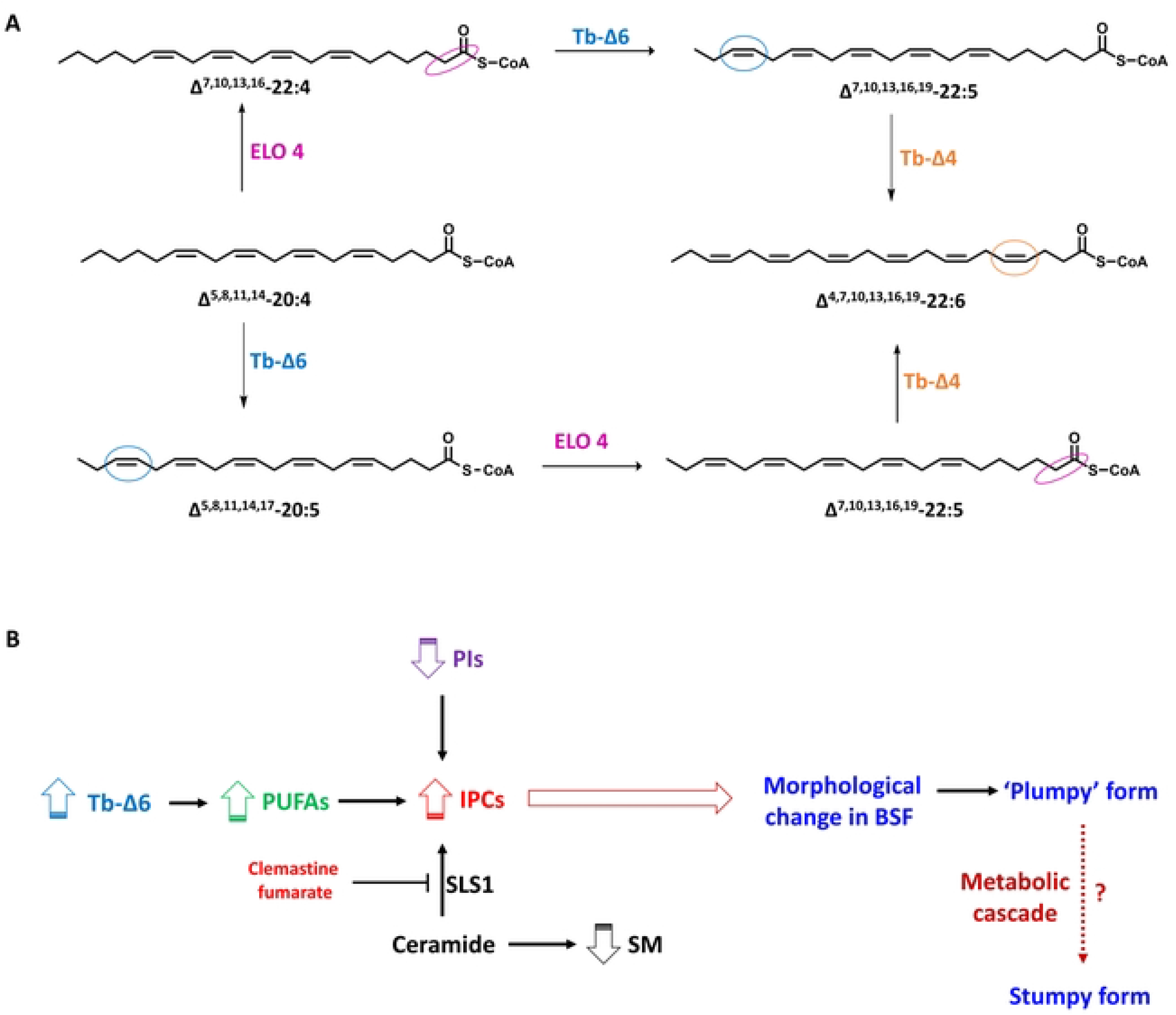
A) Biotransformation of 20C and 22C PUFAs involving Tb-Δ6. The reaction scheme depicts the biotransformation pathway for the synthesis of 20C and 22C PUFAs in *T. brucei*. The blue circles highlight the insertion of the double bond by Tb-Δ6 (blue). The orange circle highlights the insertion of the double bond by Tb-Δ4 (orange). The magenta circles highlight the elongation of two methylene groups by Tb-ELO4 (ELO 4 magenta). **B) The mechanism of production of IPC in *T. brucei* BSF overexpressing Tb-Δ6 in low-fat media.** The scheme shows the mechanism for which, in presence of higher expression level of Tb-Δ6 (light blue), Δ6-OE BSF trypanosomes produce higher level of PUFAs (green). This causes production of IPC (red) by SLS1 and a consequent reduction on PIs (purple) and SM (black), while redirecting the ceramide anchors. The synthesis of IPC by SLS1 cascade can be inhibited by clemastine fumarate. These events determine an alteration in morphology, and a shift of the cells (empty brown arrows) towards a ‘plumpy’ form. A potential cascade of metabolic events (dotted arrows) might lead the morphological changes into the stumpy pre-infective form.

Importantly, the fine regulation of the synthetic pathway of PUFAs highlighted the direct crosstalk with the lipid metabolism in *T. brucei* PCF and more evidently in BSF. (**30**) In fact, we demonstrated that PCF and BSF trypanosomes can initiate a remodelling of their FA, phospholipid and/or sphingolipid pools, as a consequence of the unbalanced levels of 20C and 22C PUFAs, ultimately causing morphological changes in the cells. The most dramatic effect of this phenomenon was revealed in BSF (Figure 8B). Surprisingly, when the expression level of Tb-ω-end-Δ4 (or Tb-Δ6) was increased, therefore when the amount of 22:5 and 22:6 was higher than normal, and upon scarce FA sources in the media, BSF trypanosomes started producing IPC (Figure 8B). This caused the consequential reduction of both SM and PI levels and redistribution of ceramide to make IPC (Figure 8B), similar to that previously observed in stumpy cells. (**30**),(**28**) These findings are very intriguing, considering that IPC is normally only produced in PCF and in the pre-adaptive to Tsetse fly stumpy stage of BSF trypanosomes, in which both life cycle satges express Tb-ω-end-Δ4 (or Tb-Δ6) at a much higher level than in BSF. (**24,29**) Taking these results into account, it seems evident that the activity of the mitochondrial IPC synthase (SLS1) (Tb927.9.9410), which is found to be upregulated in the stumpy form and PCF of *T. brucei*, might be linked to the resulting activity and product of the additional expression level of Tb-ω-end-Δ4 (or Tb-Δ6). (**29,38**) Therefore, our findings might suggest that increasing the expression of Tb-ω-end-Δ4 (or Tb-Δ6) in BSF results in the activation of SLS1 and therefore of the synthesis of IPC (Figure 8B). However, if the expression of Tb-ω-end-Δ4 (or Tb-Δ6) is turned down in PCF, this does not result in the downregulation of SLS1 and of the overall IPC synthesis, as one might expect. Therefore, *T. brucei* PCF and BSF implement different responses to the varying level of PUFAs and expression of Tb-ω-end-Δ4 (or Tb-Δ6) to control membrane fluidity. BSF parasites respond by ultimately synthesising IPC to maintain the appropriate level of membrane fluidity. PCF parasites respond by scavenging large amount FA sources from the serum, if in high-fat media, or by reducing the ratio of C22/C20 PUFAs, if in low-fat media, to maintain the membrane homeostasis and rescue their growth. We also found that, in keeping with recent studies by Machado *et al.*,(**39**) some environmental PUFAs, can have a detrimental effect on the parasite cell growth, like shown for WT and Δ6-KD PCF under DHA supplementation. Despite that, WT and Δ6-KD PCF cells showed the typical full range of cellular FA species, where an increase of C18:0/C18:1 and C18:2 ratio was observed to ensure once again membrane homeostasis and cell survival. The effect of the lipid remodelling on the cell morphology was very evident in the Δ6-OE BSF cells grown in low-fat media. Δ6-OE BSF cells shifted from an elongated slender to a ‘plumpy’ morphology, which resembles the stumpy morphology, while also expressing reasonable level of PAD1.(**32**) Once again, this is in support with is key when considering that *T. brucei* parasites encounter fat-rich and fat-poor environments, that are constantly changing going from the mammalian host to the insect vector. (**15**)

At this point a question remains open: are the overexpression of Tb-ω-end-Δ4 (or Tb-Δ6) in BSF, and the consequent production of IPC, linked through a cascade of metabolic events that might contribute to the transition from the slender BSF to the pre-adaptive stumpy form and thus to the PCF (Figure 8B)? (**29**) At this point, we can highlight that, despite the stumpy induction factor (SIF) and quorum sensing,(**8**) which are key to differentiation in *T. brucei,* cannot be and have not been induced in this study (as monomorphic *T. brucei brucei* strain 427 were used), we are still able to see some of the downstream effects (IPC synthesis, PAD1 enrichment, plumpy morphology, upregulation of Tb-Δ6) that are normally seen in the differentiation cascade of *T. brucei*. (**29**) On-going work in the lab on metabolomics and proteomics analysis is attempting to establish with more confidence whether these and other metabolic pathways that are highly active in the stumpy and PCF forms, might be up regulated in Δ6-OE BSF mutants grown in low-fat media in monomorphic and polymorphic strains. This could give a potential clearer indication of whether Tb-ω-end-Δ4 (Tb-Δ6)) expression levels are contributing to the shift from slender to stumpy and finally PCF forms.

There are many exciting questions still to be answered, but so far we clearly showed that FAs, lipids and their metabolic enzymes are key to trypanosomes’ cell growth and to their efficient adaptation to the hosts’ lipid environments. (**40**),(**33**) We demonstrated that trypanosomes are able to take advantage of those finely tuneable mechanisms to control their cell growth, based upon the level of extracellular fat sources available and the homeostasis of their membranes to meet their cellular requirements based upon the environment. These are all mechanisms that actively contribute to the morphological and metabolic needs of *T. brucei* during the different phases of the trypanosomes’ complex life cycle (Figure 8B). (**33**),

## Materials and Methods

Unless otherwise stated, all reagents and materials were purchased from Sigma, Promega, New England Bio Labs (NEB) or VWR. SPLASH LIPIDOMIX Mass Spec Standards was purchased from Avanti Polar Lipids, Gibco Iscove’s Modified Dulbecco’s Medium (IMDM) and Gibco Foetal Bovine Serum (FBS) were purchased from Thermo Fisher. SDM-79 powder was obtained from Life Technologies.

### Cell culture

*T. brucei* PCF (strain XX) parasites were grown at pH 7.4, 28°C and under 5% CO_2_ in SDM- 79 media supplemented with either 10% or 1.25% (v/v) of heat inactivated FBS, 11.5 μM haemin and 24 mM NaHCO_3_. (**41**) PCF were kept under drug selection pressure from G418 (neomycin) (15 μg/mL) and hygromycin (50 μg/mL). (**26**) *T. brucei* PCF RNAi cell lines were grown in the presence of 1.25 μg/mL phleomycin or *T. brucei* PCF overexpression cell line in the presence of 5 μg/mL blasticidin. *T. brucei* BSF (strain) parasites were grown at pH 7.4, 37°C and under 5% CO_2_. (**3**) in HMI-11 media containing 80% (v/v) IMDM, 1.25 mM pyruvic acid, 161 µM thymidine, 50 μM bathcuperoinedisulfonic acid, 1 mM hypoxanthine, 1.5 mM L-cysteine, 58 μM 1-thioglycerol and supplemented with either 10% or 5% (v/v) heat inactivated FBS. BSF were kept under drug pressure from G418 (2.5 μg/mL). (**26**) *T. brucei* BSF RNAi cell line were grown in the presence of 2.5 μg/mL phleomycin and *T. brucei* BSF overexpression cell line in the presence of 5 μg/mL blasticidin. Addition of tetracycline to the media, when required, was at a final concentration of 1 μg/mL. Cell counting was performed using a Model-TT CASY Cell Counter and Analyser system. Cells were maintained at mid-log phase and passaged every 3 days (PCF) or every 2 days (BSF).

### Chemical supplementation of *T. brucei* PCF and BSF culture media with free fatty acids

In case of free fatty acid supplementation, the media SDM-79 supplemented with 1.25% FBS and HMI-11 supplemented with 5% FBS in the presence of the appropriate selection drugs as described above and tetracycline at a final concentration of 1 μg/mL, were supplemented with either DHA (22:6) or AA (20:4) to a final concentration in the flask of 10 μM (stock solution 1M in DMSO).

### Growth curves for *T. brucei* PCF and BSF

PCF trypanosomes were grown to mid-log phase in SDM-79 supplemented with 10% or 1.25% FBS and distributed into non-vented flasks at an equal density of 5 x 10^5^ cell/mL in the presence or absence of the appropriate concentration of selection drugs and tetracycline according to the cell line. The cells were grown for 72 h and counted every 24 h using a Model-TT CASY Cell Counter and Analyser system. BSF trypanosomes were grown to mid-log phase in HMI-11 supplemented with 10% or 5% FBS and distributed into vented flasks at an equal density of 1.5 x 10^5^ cell/mL in the presence or absence of appropriate concentration of selection drugs and tetracycline according to the cell line. The cells were grown for 48 h and counted every 12 h using a Model-TT CASY Cell Counter and Analyser system.

### Determination of chromosomal location of Tb-Δ6 in *T. brucei* BSF

The region between the gene encoding for Tb-Δ6 (Tb11.v5.0580) and the downstream gene Tb11.v5.0579 on chromosome 11 was amplified from *T. brucei brucei* strain 427 genomic DNA using KOD Hot Start DNA Polymerase (Novagen), and the following forward and reverse primers 5’- AGA GTA TCC AGC TGG GAG TC -3’ and 5’- CTA CCT TGC TCC TAT TGT GC -3’.

The region between the gene encoding for Tb-Δ6 (Tb927.10.7100) and the upstream gene Tb927.10.7110 on the chromosome 10 was amplified from *T. brucei brucei* strain 427 genomic DNA using KOD Hot Start DNA Polymerase (Novagen), and the following primers 5’- CTA CCT TGC TCC TAT TG TGC -3’ and 5’- CTA TAC GTT GCG GTG ATG G -3’.

### Generation of Tb-Δ6 RNAi construct using p2T7-177 vector

The ORF of Tb-Δ6 (Tb11.v5.0580.1) gene was amplified from *T. brucei brucei* strain 427 genomic DNA using KOD Hot Start DNA Polymerase (Novagen), and the following forward and reverse primers 5’-TTT T<u>TG GAT CC</u>A TGA GTT CGG TAA AGA GTA AAG G-3’and 5’-TTT T<u>TC TCG AG</u>C AAC CGT TTG TCT TCT ATG C-3’ containing *BamH*I and *Xho*I restriction sites (underlined regions). The purified product digested with *BamH*I and *Xho*I and ligated into appropriately digested p2T7-177, containing the phleomycin (Phleo) resistance cassette. The plasmid obtained was defined as p2T7-177-Tb-Δ6-Phleo and the presence of the insert was confirmed by PCR colony screening using with the following sets of forward and reverse primers 5’-ATAGAGATCTAGCCGCGGTGG–3’ and 5’– CATGATGAATGATCCAACTGATGC–3’ (expected product size 630 bp), and 5’- CCCTATCAGTGATAGAGATCTCC-3’ and 5’–CAAACGAACCTCATCATGACTTGG–3’ (expected product size 702 bp), using GoTaq DNA polymerase (Promega, and sequencing at Eurofins Genomic.

### Generation of Tb-Δ6 overexpression construct using pLew-100-C-term-6HA vector

The ORF of Tb-Δ6 (Tb11.v5.0580.1) gene was amplified from *T. brucei brucei* strain 427 genomic DNA using KOD Hot Start DNA Polymerase, using the following forward and reverse primers 5’-ATA GTA C<u>AA GCT T</u>AT GAG TTC GGT AAA GAG TAA AGG -3’ and 5’- GAT AGC <u>TTA ATT AA</u>C AAC CGT TTG TCT TCT ATG C-3’ containing *Hind*III and *Pac*I restriction sites (underlined regions). The purified product was digested with *Hind*III and *Pac*I and ligated into appropriately digested pLew-100-C-term-6HA, containing the blasticidin (BSD) resistance cassette. The plasmid obtained was defined as pLew-100-Tb-Δ6-C-term- 6HA-BSD and the presence of the insert was confirmed by was confirmed by PCR colony screening (expected size product 1499 bp), with the following forward 5’-CTC GTC CCG GGC TGC ACG CGC CTTCCG-3’ and reverse 5’-CCT GCA GGC GCA CCT CCC TGC TGT G-3’ primers, using GoTaq DNA polymerase (Promega), and sequencing at Eurofins Genomic.

### Transfection of *T. brucei* PCF and BSF

1 x 10^7^ PCF and BSF mid-log phase cells were harvested and transfected using Lonza T-cell nucleofector kit and Amaxa-electroporator using program X-014 (PCF) and X-001 (BSF) as reported previously (**42**). The transfected trypanosomes were transferred into a 24 well/plate and incubated at 28°C (PCF) or 37°C (BSF) under 5% CO_2_ for 24 h. At this point, 2.5 μg/mL (PCF) or 5 μg/mL (BSF) of phleomycin, and 10 μg/mL (PCF and BSF) of blasticidin were added to the flasks’a. After several days, the cells that showed the highest growth rate were selected and OE-Δ6 BSF and PCF, KD-Δ6 BSF and PCF and OK-Δ6 PCF genotypes were verified by PCR analysis of genomic DNA isolated from parental and RNAi (5’-TTT TT<u>G</u> <u>GAT CC</u>A TGA GTT CGG TAA AGA GTA AAG G-3’ and 5’-ATG GCC AAG TTG ACC AGT GC-3’), overexpression (primers used 5’-GTC TCA AGA AGA ATC CAC CCT C-3’and 5’- GAT AGC <u>TTA ATT AA</u>C AAC CGT TTG TCT TCT ATG C-3’) and add-back (both pairs of primers listed above) cell lines.

### Total RNA extraction and qRT-PCR

Total RNA was isolated from 1x10^8^ mid-log BSF and PCF *T. brucei* cells using the RNeasy mini kit (Qiagen). RNA was treated with Precision DNase Kits (Primer Design) to remove any excess of gDNA carried over from the extraction of total RNA. A *Tb*Δ6-specific cDNA was generated and amplified using a mini-exon-specific forward primer 5’-ACG CTA TTA TTA GAA CAG TTT CTG TAC TAT ATT GAC TTT C AT GAG TTC-3’, in combination with an ORF-specific reverse primer 5’-TGT AGT GTC GGG TTC ATC GG-3’, specific for the sequence of *Tb*Δ6.(**43**) The *T. brucei* actin gene was used as endogenous control (*Tb*Act) using a mini-exon-specific forward primer 5’-GCG GAG ACT TCG AAC GCT ATT ATT AGA ACA GTT TCT GTA -3’ and ORF-specific reverse primer 5’-CTT CAT GAG ATA TTC CGT CAG GTC-3’, specific for the sequence of *Tb*Actin.(**43**) cDNA synthesis and qRT-PCR were performed in a one-pot reaction using Luna® Universal One-Step RT-qPCR Kit (NEB). A QuantStudio Real-Time PCR System (ThermoFisher) thermocycler was used following standard cycling conditions. The data were analysed by the 2^−ΔΔCT^ method normalising with *Tb*Actin.

### Lipid extraction

2 x 10^7^ cells of mid log phase *T. brucei* PCF and BSF (̴were collected by centrifugation at 800 x *g* for 10 min at room temperature. The cell was washed with either phosphate-buffered saline (PBS) (for PCF) or trypanosome dilution buffer (TDB) (5 mM KCl, 80 mM NaCl, 1 mM MgSO_4_, 20 mM Na_2_HPO_4_, 2 mM NaH_2_PO_4_, and 20 mM glucose at pH 7.7) (for BSF), after which they were re-suspended in 100 μL of PBS and transferred to a glass vial. 375 μL 2:1 (v/v) of MeOH:CHCl_3_ solution was added for biphasic separation based on the following method described by Bligh-Dyer.(**44**) The samples were vigorously agitated at 4°C overnight. The internal standards Avanti-SPLASH were prepared by transferring the 1mL MeOH solution in the vial and suspending in 1 mL 1:1 (v/v) of MeOH:CHCl_3_. 20 μL of this were added to the samples at the point of extraction. Samples were made biphasic by the addition of 125 μL CHCl_3_ and 125 μL distilled sterile water, followed by vigorous agitation after each addition. The samples were centrifuged at 1000 x *g* for 5 min to allow the aqueous and the organic layers to separate. The organic phase was transferred into a new glass vial and dried under nitrogen gas stream to obtain the total lipid extract ready to use and stored at 4°C until further analysis.

### Electrospray ionization mass spectrometry and tandem mass spectrometry (ESI-MS/MS)

Lipid samples were analysed by electrospray ionization tandem MS (ESI-MS/MS) with an AB- Sciex Qtrap 4000 triple quadrupole mass spectrometer fitted with an Advion TriVersa NAnoMAte nanoelectrospray. Survey scans in negative mode were used for the detection of PE, EPC, PI, IPC, PS, PG and PA (cone voltage = 1.25 kV). Positive ion mode survey scans were used to detect PC and SM (cone voltage = 1.25 kV). In negative ion mode, a precursor of m/z 196 scans were used to detect PE and EPC species, a precursor of m/z 241 scans for IPC and PI species, a precursor of m/z 153 scans for PG and PA, (collision energy, CE 60 eV). In positive ion mode, a precursor of m/z 184 scans were used to detect PC and SM species (CE 60 eV). Neutral loss of a precursor of m/z 87 scans were used for PS (CE 60 eV). Spectra were acquired over or within a range of 120-1000 m/z, and each spectrum represents a minimum of 30 consecutive scans with nitrogen collision gas. Samples were run using a 1:1 solvent mixture of 2:1 (v/v) MeOH:CHCl_3_ and 6:7:2 (v/v) acetonitrile:isopropanol:dH_2_O. Individual lipid species were annotated according to their acyl composition determined by daughter ion scans produced and compared to previous lipid identification by Richmond et al.,(**28**) and to theoretical values contained in the Lipid Metabolites and Pathways Strategy consortium database (LIPID MAPS, http://www.lipidmaps.org/).

### Fatty acids transmethylation and gas chromatography – Mass spectroscopy (GC-MS) samples preparation

Transmethylation of the fatty acids was performed on the dry total lipid extracts.(**45**) The reaction (total volume 1 mL) was conducted in a glass vial. 100 μL of toluene were added, followed by 750 μL of MeOH and 150 μL of 8% solution of HCl in MeOH:H_2_O (85:15 (v/v)) to allow the formation of fatty acid methyl esters (FAMEs). The reaction was left to react at 45°C overnight. Upon drying under a nitrogen gas stream, the FAMEs were extracted with a 1:1 (v/v) hexane:H_2_O solution. The FAME extracts were dried under a nitrogen gas stream and dissolved in dichloromethane, typically 15 μL, of which 1-3 μL were analysed by GC-MS on an Agilent Technologies GC-6890N gas chromatograph coupled to an MS detector-5973. Separation by GC was performed using a Phenomenex ZB-5 column (30 m x 25 mm x 25 mm), with a temperature program of 70°C for 10 min, followed by a gradient to 220°C, at 5°C/min, which was maintained at 220°C for a further 15 min. Mass spectra were acquired from 50-500 amu.(**46**) FAME species were assigned by comparison of the retention time, fragmentation pattern, use of FAME standards, and the online FAMEs mass spectrometry database (http://www.lipidhome.co.uk/ms/methesters/me-arch/index.htm).

### Western blot probed with an anti-HA antibody and samples preparation

1 x 10^7^ mid-log-phase PCF and BSF parasites were harvested by centrifugation at 800 x g for 10 min. The cells were washed in PBS twice and re-suspended in 50 μL of 2 x SDS sample buffer and incubated at 95°C for 10 min. SDS-PAGE was performed on 12% acrylamide gel.

The protein gel was then transferred onto a membrane (GE Healthcare Life Science UK) and the membrane blocked in 5% semi skimmed milk powder in PBS. The membrane was incubated for 1 h at room temperature with Odyssey blocking buffer and 0.2 % PBS-T (PBS containing 0.2 % (v/v) Tween-20) with rat anti-HA clone 3F primary antibody (Clontech) at 1:1,000 dilution. The membrane was washed for 3 x 10 min with 0.2 % PBS-T buffer. Subsequently, the membrane was incubated for 1 h at room temperature with secondary goat anti-rat conjugated with DyLight 800 (Licor IRDye 800) antibody at 1:10,000 dilution. The membrane was washed as above, before the fluorescence was detected with an Odyssey western blot detection system (Licor Odyssey).

### Immunofluorescence microscopy for *T. brucei* PCF and BSF

1 x 10^6^ PCF and BSF trypanosomes were harvested at 800 x g for 10 min and washed with PBS. (i) For mitochondrial visualisation, cells were incubated for 30 min at 28°C or 37°C, for PCF and BSF respectively, with 100 nM of MitoTracker red CMZRos in 100 μL of fresh SDM- 79 or HMI-11. The cells were spun at 3,000 rpm for 2 min and re-suspended in 100 μL of fresh media and incubated at 28°C or 37°C for 15 min, to allow the MitoTracker to enter the mitochondria. The cells were spun again as above and washed with PBS. The cells were resuspended in 100 μL of 4% paraformaldehyde (PFA) in PBS and incubated at room temperature for 15 min. After this time, the cells were spun at 3000 rpm for 2 min, the supernatant removed, and the cells washed with PBS and re-suspended in 100 μL fresh PBS. 50 μL of cells were added to Polysine® slides (VWR 631-0107) and allowed to adhere overnight at room temperature in a sealed container on a damp tissue to prevent evaporation. The cells were re-hydrated with 100 μL of dH_2_O for 5 min and washed with 100 μL of PBS for 2 x 5 min using a pipette. The cells on the slide were incubated with 100 mM glycine in PBS for 5 min. The cells were washed for 3 x 5 min with 100 μL of PBS. At this point 50 μL of 0.1% TritonX-100 solution in PBS were added to permeabilise the cells for 10 min. The cells were washed for 3 x 5 min with 100 μL of PBS and blocked for 20 min with 1% BSA in PBS. The cells were washed for 1 x 5 min with 100 μL of PBS. (ii) For HA-tag localisation, the primary rat anti-HA antibody was diluted to 1:500 in 1% BSA-PBS, and the cells were incubated with this solution at room temperature for 2 h. The cells were washed with 100 μL of 1% BSA-PBS for 3 x 5 min. The secondary chicken anti-rat Alexa Fluor 594 antibody (Thermo Fisher) was diluted to 1:1,000 in 1% BSA-PBS, and the cells were incubated with this solution at room temperature for 2 h. The slides were washed for 3 x 5 min with PBS in a Coplin jar. The slides were thoroughly dried with the use of a pipette. (iii) For Tb-PAD1 visualisation, the cells were blocked for 45 min with 2% BSA in PBS in a humidity chamber at 37°C. The cells were washed for 1 x 5 min with 100 μL of PBS. Primary rabbit anti-α-PAD1 antibody (kindly given by Prof Keith Matthews, The University of Edinburgh) was diluted to 1:1,000 in 2% BSA-PBS, and the cells were incubated with this solution at 37°C in a humidity chamber for 45 min. The cells were washed with 100 μL of 2% BSA-PBS for 3 x 5 min. The secondary chicken anti-rabbit Alexa Fluor 488 antibody (Thermo Fisher) was diluted to 1:5,000 in 2% BSA-PBS, and the cells were incubated with this solution at 37°C in a humidity chamber for 45 min. The slides were washed for 3 x 5 min with PBS in a Coplin jar. (iv)To stain the DNA, the cells were incubated with 50 μL DAPI (4,6-diamidino-2-phenylindole) (2 μg/mL, in PBS) for 5 min in the dark. The slides were washed for 3 x 5 min with PBS. A drop of antifade agent was added to each sample on the slide, covered with a coverslip and let dry at room temperature overnight, before sealing the coverslips with nail polish. Images were acquired with a fluorescence confocal microscope (DeltaVision Imaging System), connected to a digital camera system, and processed by softWoRx 1.3 image analysis software.

### Live/dead dose-response cytotoxicity assay in *T. brucei* BSF for clemastine fumarate

The Alamar Blue cell viability assay was performed by incubating 1 × 10^3^ cells/well (final volume 200 μL) mid-log-phase *T. brucei* BSF cells with the clemastine fumarate (DMSO stock of 50 mM) and 2-fold serially diluted in HMI-11 in a concentration range from 200-0.20 μM) at 37°C with 5% v/v CO_2_ for 72 h. After this, 10 μL Alamar Blue (1.1 mg/mL resazurin sodium salt in PBS) was added to all wells and the cells incubated for a further 7 h before recording fluorescence.(**47**) Fluorescence was recorded using a FLx 800 plate reader (BioTek) with excitation wavelength 530–535 nm and emission wavelength at 590–610 nm and data were processed using Gen5 Reader Control 2.0 Software (BioTek). EC_50_ values were determined using a 4-parameter non-linear logistic regression equation using GraFit 5.0 (Erithacus Software). SD values were calculated based on curve fitting to n biological replicates performed in parallel.

### DMDS-FAME adducts synthesis from whole cell lipid extract of *T. brucei* PCF and BSF

FAMEs, obtained after transmethylation of lipid extracts from 1 x 10^7^ cells of *T. brucei* BSF and PCF cells were dissolved in 80 μL of hexane prior to addition of 100 μL DMDS. A catalytic amount of iodine solution in diethyl ether (60 mg/mL) was added to the reaction mixture under agitation for 12 h at room temperature.(**48**) The reaction was then added with 2 mL hexane and 2 mL 0.1 N sodium thiosulfate solution and vigorously agitated until the solution discoloured completely. (**48**) The top organic layer was extracted and dried under a nitrogen stream. The final DMDS-FAME adducts mixture was dissolved in 15 μL DCM and analysed by GC-MS as described above.

### Statistical analysis

For all growth curves, FAMEs and lipids experiments values are the mean of three independent biological replicates (n=3). Error bars represent the standard deviation of each mean calculated by GraphPad PRISM 6.0. Statistical analysis, were required, was also performed by GraphPad PRISM 6.0 using One-way or 2-ways ANOVA multiple comparisons or based on a Tukey or Dunnet t-test or unpaired t-test with a 95% confidence interval.

## Acknowledgement

We would like to thank the Engineering and Physical Sciences Research Council, University of St. Andrews, and the EPSRC Centre for Doctoral Training in Critical Resource Catalysis (CRITICAT) for financial support [Ph.D. studentship to MC; Grant code: EP/L016419/1].

## Abbreviations

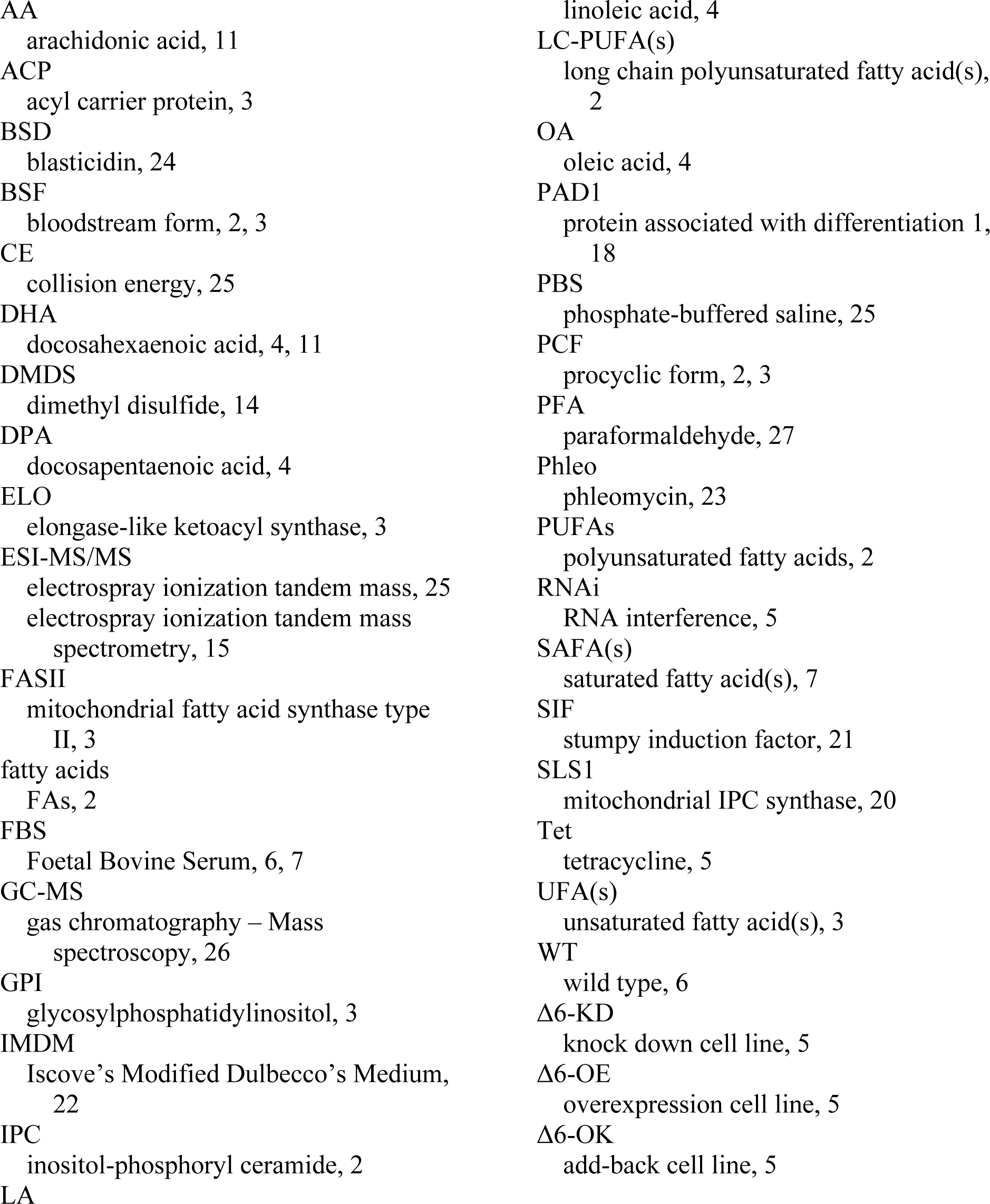

## Supplementary Information

Appendix A FAME FBS-FAs

Appendix B Tbrucei BSF-FAs and Lipid Content Appendix C Tbrucei PCF-FAs and Lipid Content Appendix D. Tb-D6-qRT-PCR.xlsx

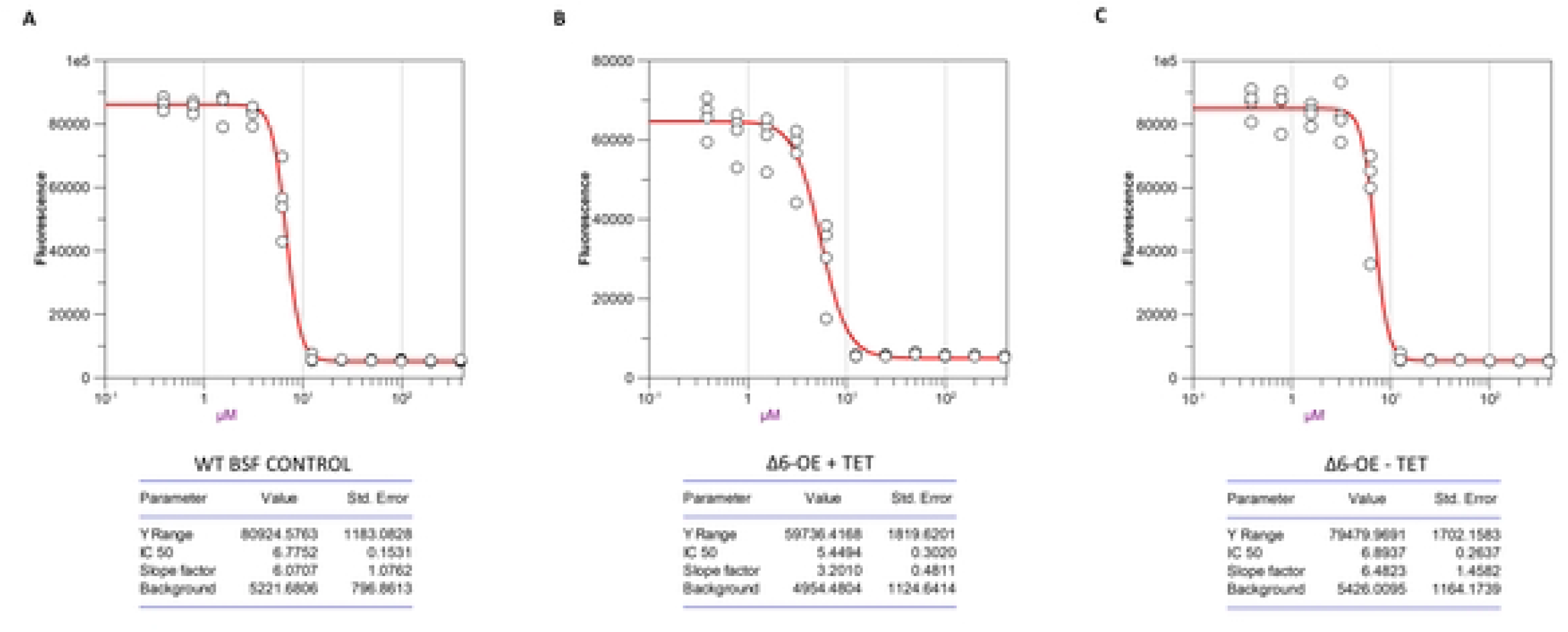

